# Identification of novel alternative splicing events associated with tumorigenesis, protein modification, and immune microenvironment in early-onset gastric cancer

**DOI:** 10.1101/2020.08.22.262717

**Authors:** Jian Zhang, Ajay Goel, Lin Zhu

**Affiliations:** Department of Pharmaceutical Sciences, Irma Lerma Rangel College of Pharmacy, Texas A&M University, College Station, TX, USA; Beckman Research Institute, City of Hope Comprehensive Cancer Center, Biomedical Research Center, Monrovia, CA, USA

**Keywords:** alternative splicing, immune microenvironment, early-onset gastric cancer, protein phosphorylation, protein glycosylation, alternative polyadenylation

## Abstract

**Background:** Alternative splicing (AS), e.g. tandem alternative polyadenylation (TAPA), has emerged as major post-transcriptional modification events in human disease. However, the roles of AS and TAPA in early-onset gastric cancer (EOGC) have not been revealed.

**Methods:** The global AS profiles of 80 EOGC patient samples from the European Nucleotide Archive (PRJNA508414) were analyzed. The EOGC-specific AS events (ESASs) were identified in both EOGC and adjacent non-tumor tissues. Functional enrichment analysis, Splicing network, Alternative Polyadenylation (APA) core factor network, and cell abundancy analysis were performed. Furthermore, the landscapes of AS events in the varied subtypes of EOGC patients, including various protein modifications and viral infections, were evaluated.

**Results:** Overall, 66,075 AS events and 267 ESASs were identified in EOGC. In these events, 4809 genes and 6152 gene isoforms were found to be aberrantly expressed in EOGC. The Gene Ontology (GO) and Kyoto Encyclopedia of Gene and Genome (KEGG) pathway analyses showed that significant pathway alterations might exist in these AS events, genes, and gene isoforms. Moreover, the Protein-protein interaction (PPI) network analysis revealed that UBC, NEK2, EPHB2, and DCTN1 genes were the hub genes in the AS events in EOGC. The immune cell infiltration analysis indicated a correlation between the AS events and the cancer immune microenvironment. The distribution of AS events in varied EOGC subtypes was uneven. The numbers of AS events related to protein phosphorylation and glycosylation were 82 and 85, respectively, which suggested a high association between AS events and protein modification in EOGC.

**Conclusion:** The study highlighted the vital roles of AS in EOGC, including modulating the specific protein modification and reshaping the cancer immune microenvironment, and yielded new insights into the diagnosis of EOGC as well as cancer treatment.

## Introduction

Gastric cancer (GC), a morbid and frequently lethal malignancy, is one of the most common cancer and the leading cause of cancer death worldwide, especially in east Asia [1]. For most patients, GC is usually associated with unfavorable prognoses and can be only diagnosed at the relatively late stages, resulting in limited treatment options. The major types of cancer, including the brain, cervix, esophageal squamous cell carcinoma, Kaposi sarcoma, larynx, lung, and non-Hodgkin lymphoma, remain a relatively low incidence rate among young adults. However, the cases of gastric non-cardia cancer in young adults kept increasing from 1995 to 2014 in US [2]. The GC occurring in the patients under 45 years old is defined as EOGC [3]. Though the genetic and environmental factors have been identified to be associated with GC, the occurrence of EOGC remains largely unexplained.

Accumulating evidence showed that somatic CDH1 or TGFBR1 gene mutation and proteogenomic alternation were remarkable in younger adults, which may promptly unleash the GC [4, 5], suggesting the importance of the post-transcriptional regulation in EOGC. AS, one of the key post-transcriptional events, can generate various mRNA transcripts (isoforms) and affect their stability as well. As a result, the downstream protein variants translated from these mRNA isoforms vary in their sequences [6]. On the one hand, the protein synthesis is directed by the human genome, but on the other hand, the protein diversity is ensured to satisfy various biological processes [7]. Two studies have profiled the AS events in The Cancer Genome Atlas (TCGA) gastric carcinomas [8] or in Epstein-Barr virus (EBV) associated gastric carcinomas [9], which ascertained the roles of AS events in gastric cancer. Li et al. found the novel AS events and peptides during mouse stomach formation [10]. Furthermore, a comparative genomic study has been performed and identified a distinct molecular expression profile in the EOGC rather than late onset gastric cancer (LOGC) [11]. All these studies suggest that EOGC may have a varied AS event landscape compared to other types of GCs.

The TAPA event introduces two or more poly(A) sites to the 3’ UTR of a gene which is distributed across a wide variety of cancers and modulates the sensitivity of certain anticancer drugs [12, 13]. The shift of TAPA events shows a cancer-specific and cell organelle-specific mode [14]. It was reported that the APA-guided shortening of NET1 gene 3’ UTR enhanced mRNA transcriptional activity and promoted the metastasis of gastric cancer cells [15]. Some studies defined TAPA as a subtype of AS, while others consider it a pre-mRNA processing [16]. In this study, TAPA was analyzed as AS.

Unfortunately, so far, only a few genome-wide association studies on EOGC have been performed and none of them have taken into account the effects of AS events on EOGC. To our knowledge, the roles of AS in EOGC remains vogue. Herein, the major aims of this study are to analyze the AS events including TAPA events and evaluate their roles in EOGC. The study contains the large-scale RNA sequencing data generated from EOGC samples. The human transcriptome is surveyed to identify the genome-wide AS events in EOGC and the ESAS are identified thereafter. Furthermore, the biological functions of these events are explored. The AS events in different EOGC subtypes are deciphered and their association with the molecular features and tumor immune microenvironment are profiled.

## Methods

### Data source

The RNA-Seq data of an independent cohort (tumor tissues and adjacent normal tissues from 80 EOGC patients) was retrieved from the European Nucleotide Archive (study accession: PRJNA508414). One adjacent normal tissue sample (SRR8281377) was excluded from this dataset due to the file error. These patients were histologically diagnosed as GC and with the age at the time of surgery ≤ 45 years.

### RNA-Seq analysis of AS events

Paired end reads per sample data were generated using the HiSeq 2500 platform sequencer (Illumina). Alignment was performed using Whippet (version 0.11) [17] to the Homo sapiens GRCh37.75 genome assembly (ftp://ftp.ensembl.org/pub/release-75//fasta/homo_sapiens/dna/Homo_sapiens.GRCh37.75.dna.primary_assembly.fa.gz). Beside default settings, Whippet was run with the “biascorrect” option to implement GC-content and 5’ sequence bias correction. PSI value for AS events as well as read count of genes and gene isoforms were generated by Whippet. The output file (diff. format) for comparative analysis of PSI was generated by Whippet-delta.jl using default parameters. The comparative analysis for genes and isoforms was computed by limma [18] and/or edgeR [19] package.

The Percent Spliced-in Index (PSI or Ψ; range from 0.0 to 1.0) value was defined as the proportional abundance of certain AS events and was calculated for the eight types of AS events. The AS events were determined using maximum likelihood estimation by the expectation-maximization (EM) algorithm. To generate more accurate AS event profiling, the stringent filters (percentage of samples with PSI values > 95%, minimum PSI value > 0.05) were implemented. For gene and gene isoform profiles, genes or gene isoforms with minimum Counts Per Million (CPM) > 0.5 and percentage of samples with read count values > 95% were included.

### Identification and pathway analysis of ESAS, genes, and gene isoforms

To filter ESAS in gastric cancer, the PSI values of AS events were compared between the tumor and the matched adjacent normal tissues. AS events with missing data exceeding 90% of all subjects were excluded. Statistical analysis was performed in the differentially expressed AS events with > 0.9 probability and absolute fold change ⩾ 0.1. The genes or gene isoforms with absolute fold change ⩾ 1 and false discovery rate (FDR) < 0.05 were defined as EOGC-specific genes and gene isoforms. Interactive sets among the eight types of AS were visualized by UpSetR (version 1.4.4) [20].

The associations between the parent genes of ESAS and biological annotation terms (Gene Ontology [GO] and Kyoto Encyclopedia of Gene and Genome [KEGG] pathway) were detected using the clusterProfiler package [21]. NetworkAnalyst updated on 05/2020 was applied to analyze the parent genes of each ESAS for constructing the visualized PPI network [22]. The STRING Interactome was selected in the PPI database (confidence score cut-off value 900).

### Immune cell infiltration and ESAS event analysis

The differential gene expression between the tumor and normal tissues and the percentage of cell abundance in the TCGA STAD cohort were calculated by TIMER2 [23]. The gene expression data from our study was input into ImmuCellAI [24] to impute the abundance of 24 types of immune cells in EOGC. Spearman’s rank correlation analysis was used to calculate the correlations between the PSI values of ESASs and immune cell type. The threshold of Spearman’s rank correlation coefficients was set to > 0.4 or < −0.4, and the BH adjusted p-value < 0.05.

### Construction of correlation network among splicing factors, APA core factors, and ESAS events

From the database [25] and publication [26], we selected 71 experimentally validated human splicing factors and 22 APA core factors to build the splicing factor correlation network and APA core factor correlation network. Spearman’s rank correlation analysis was used to impute the correlations between the expression of splicing factors or APA core factors and PSI values of ESASs or TAPAs. The threshold of Spearman’s rank correlation coefficients was set to > 0.4 or < −0.4, and the BH adjusted p-value < 0.05. The correlation network was constructed and visualized by Cytoscape (version 3.7) [27]. All statistical analyses were performed by the R language [28] and p-value < 0.05 was considered statistically significant unless specified.

### Identification of EOGC subtype relevant AS events

Data of the EOGC EBV status, microsatellite instability (MSI) status, protein phosphorylation subtype, and protein glycosylation subtype were accessed from Mun et al. [5]. AS events which have > 0.9 probability of being differentially expressed between different EBV status, MSI status, protein phosphorylation subtypes, and protein glycosylation subtypes were defined as the subtype relevant AS events in EOGC. The groups with the sample numbers of less than four were not included in the analysis. The PSI difference between groups was required to be larger or equal 0.1 for at least two groups to ensuring biological significance. The analysis was performed for each subtype separately.

## Results

### Overview of AS events in EOGC cohort

The corresponding RNA-Seq data of 80 EOGC patients were used to establish the integrated AS event profiling. We identified 66,075 AS events from 11,282 genes, which accounted for 53.0% of non-redundant human protein-coding genes [29]. Beside two special patterns of AS events: tandem transcription start site (TSS) and tandem alternative polyadenylation Site (TAPA), these AS events were classified into six canonical splicing patterns: core exon (CE), alternative acceptor splice site (AA), alternative donor splice site (AD), retained intron (RI), alternative first exon (AF), and alternative last exon (AL), as illustrated in Figure 1a. Among these splicing patterns, TAPA occurred most frequently (71.0%) (Figure 1b).

**Figure 1.**
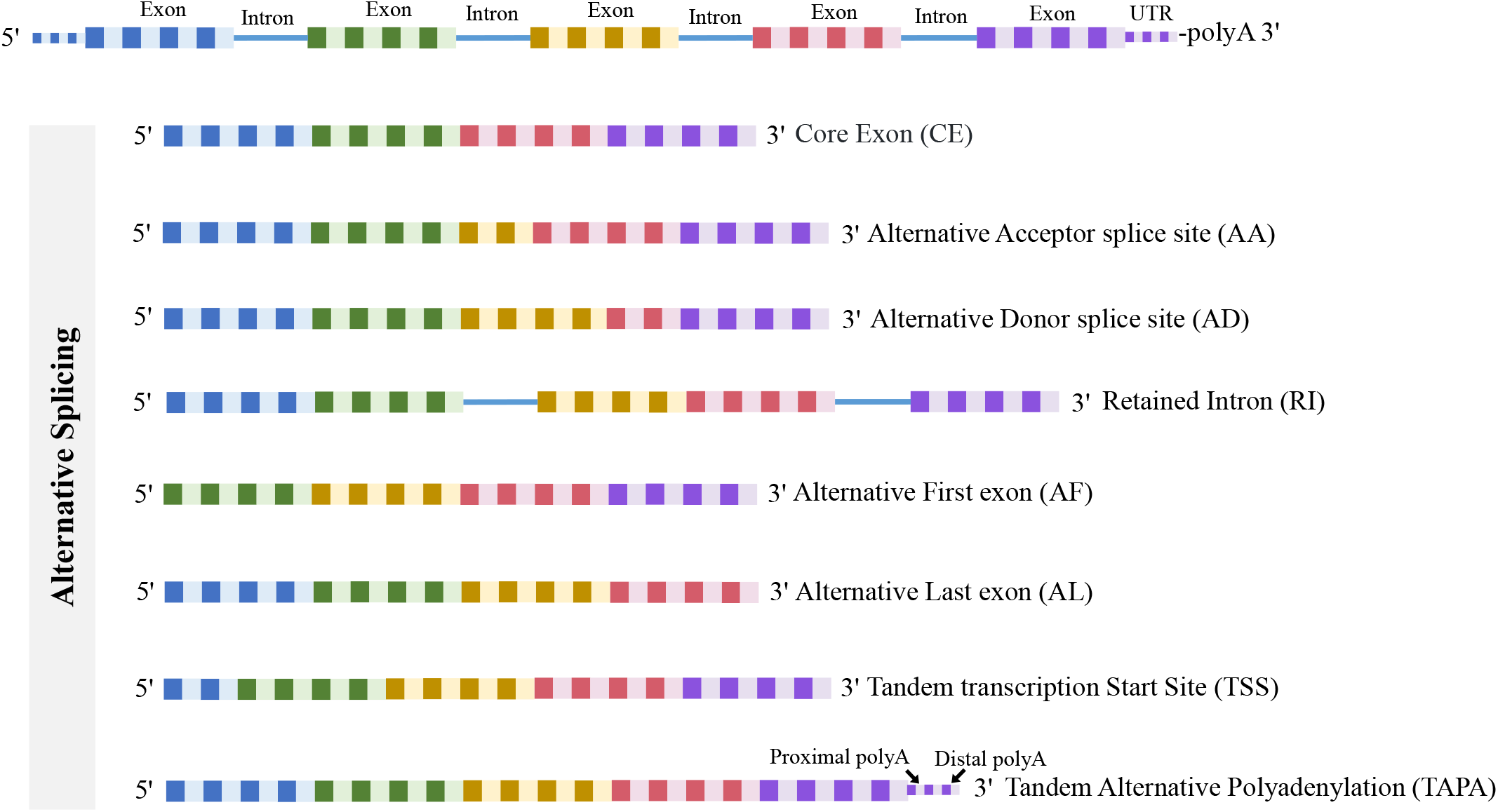
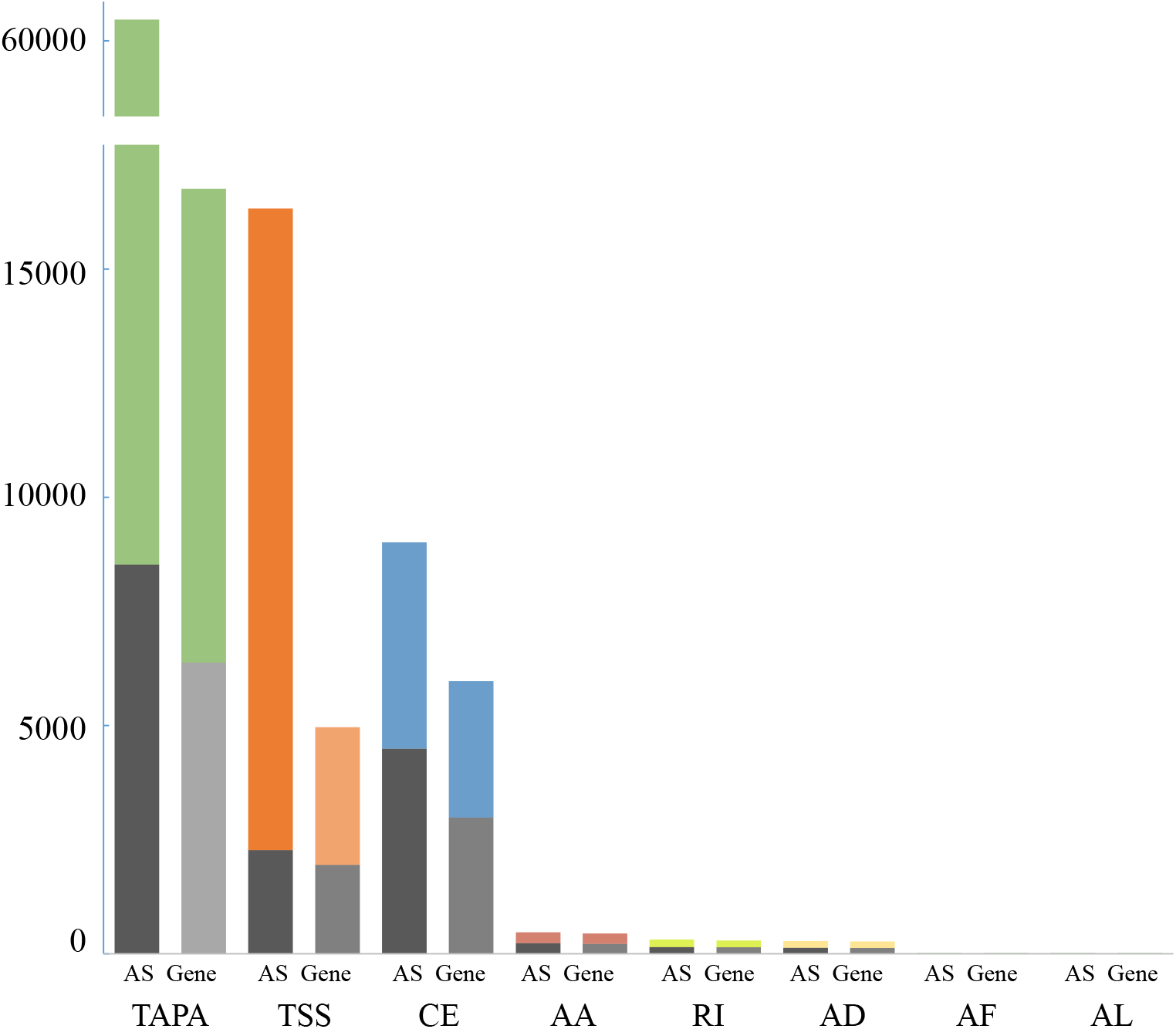
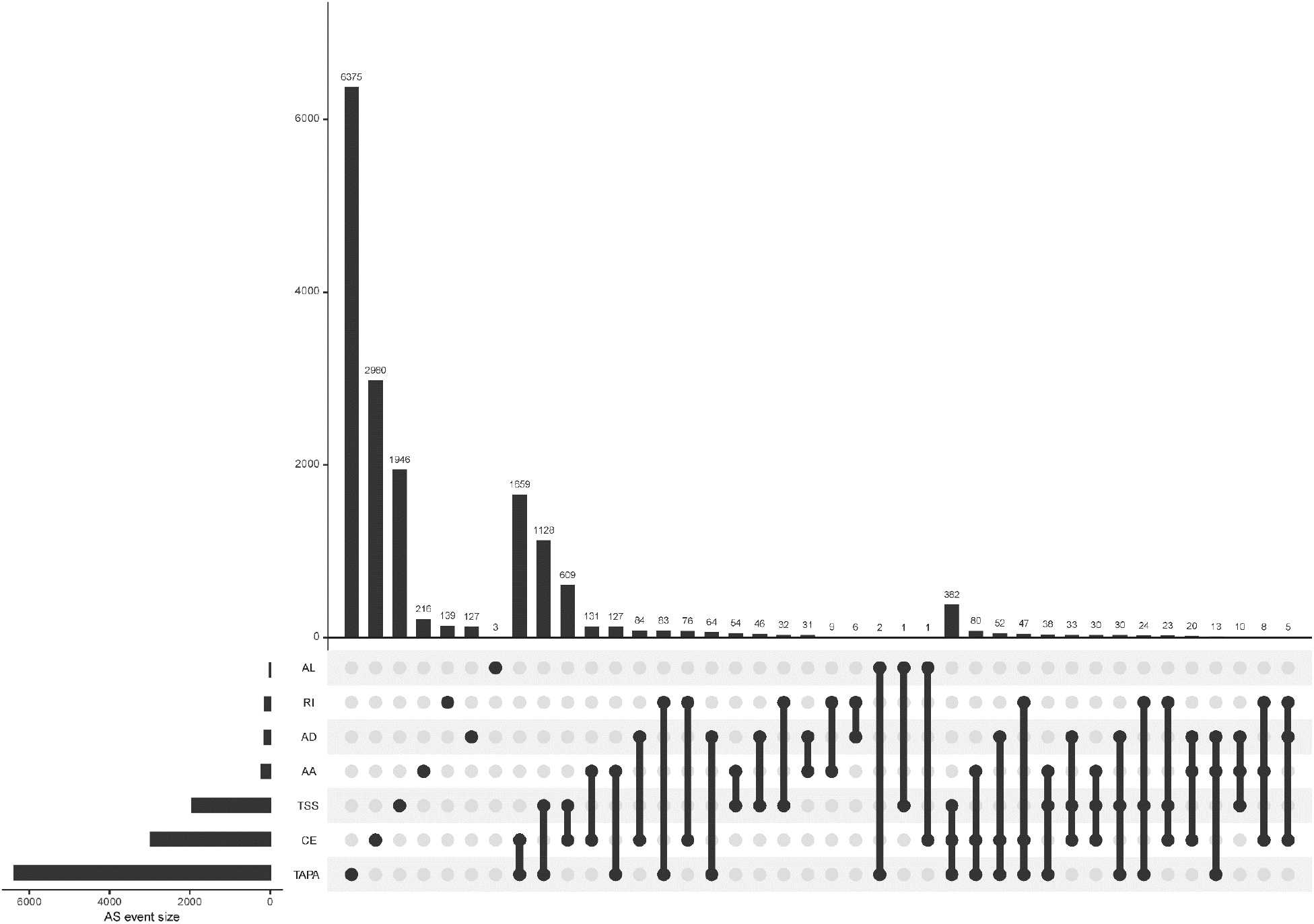
The profiling of AS events in EOGC. (a) Scheme of six canonical and two novel patterns of AS events: CE, AA, AD, RI, AF, and AL, TSS and TAPA; (b) Number of AS events and their parent genes in EOGC patients. Bar color represents the filtered AS events and their parent genes. Black bars represent the AS events and their parent genes filtered using stringent filters; (c) Interactive sets among seven patterns of AS events (n = 15,803) shown in an UpSet plot.

Some gene isoforms showed very low redundancy, thus, we screened the AS events with a series of filters (percentage of samples with PSI values ⩾ 75, PSI value ⩾ 0.05). Consequently, a total of 15,803 AS events from 8,359 genes were obtained. After filtering, AF events were excluded. The analysis indicated that TAPA was still the top AS pattern (54.0%) (Figure 1c). After removal of the duplicated genes in one AS pattern, the Upset plots were created to quantitatively analyze interactive sets of the remaining seven patterns of AS events. As shown in Figure 1c, most of the genes had more than one pattern of AS events, among which some genes had up to three different splicing patterns.

### Identification of ESAS events

To identify the EOGC-specific AS events, we compared the PSI values between 80 paired tumor tissues and 79 adjacent normal tissues. A total of 267 EOGC-specific AS events (ESASs) from 228 genes were identified (Figure 2a, Table S1). Although a large number of AS events were detected in the EOGC cohort, a relatively small proportion of AS events were identified as ESAS. TSS and TAPA events accounted for 36.3% and 24.0% of ESAS, respectively (Figure2b). The uneven distribution of AS patterns in tumor and adjacent normal tissues indicated that they might play important roles in early-onset gastric tumorigenesis.

**Figure 2.**
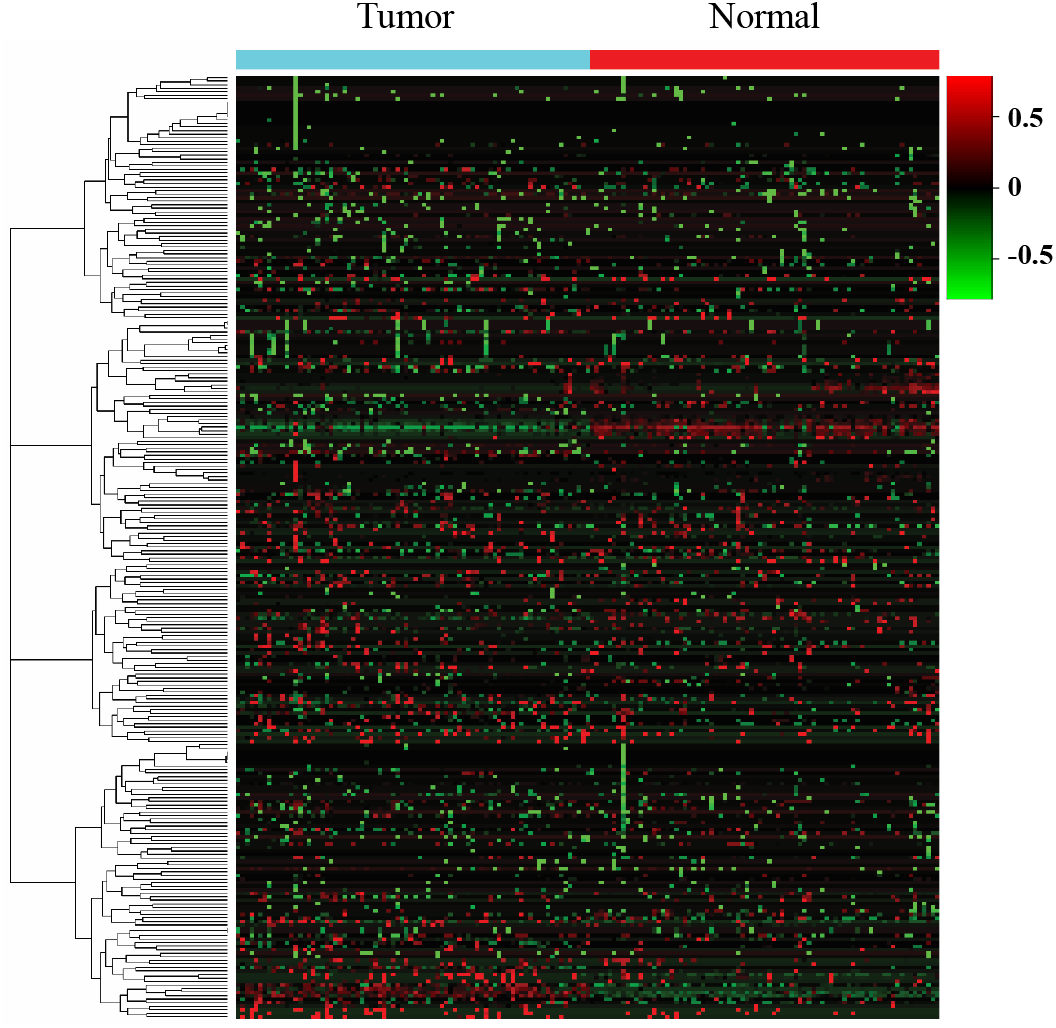
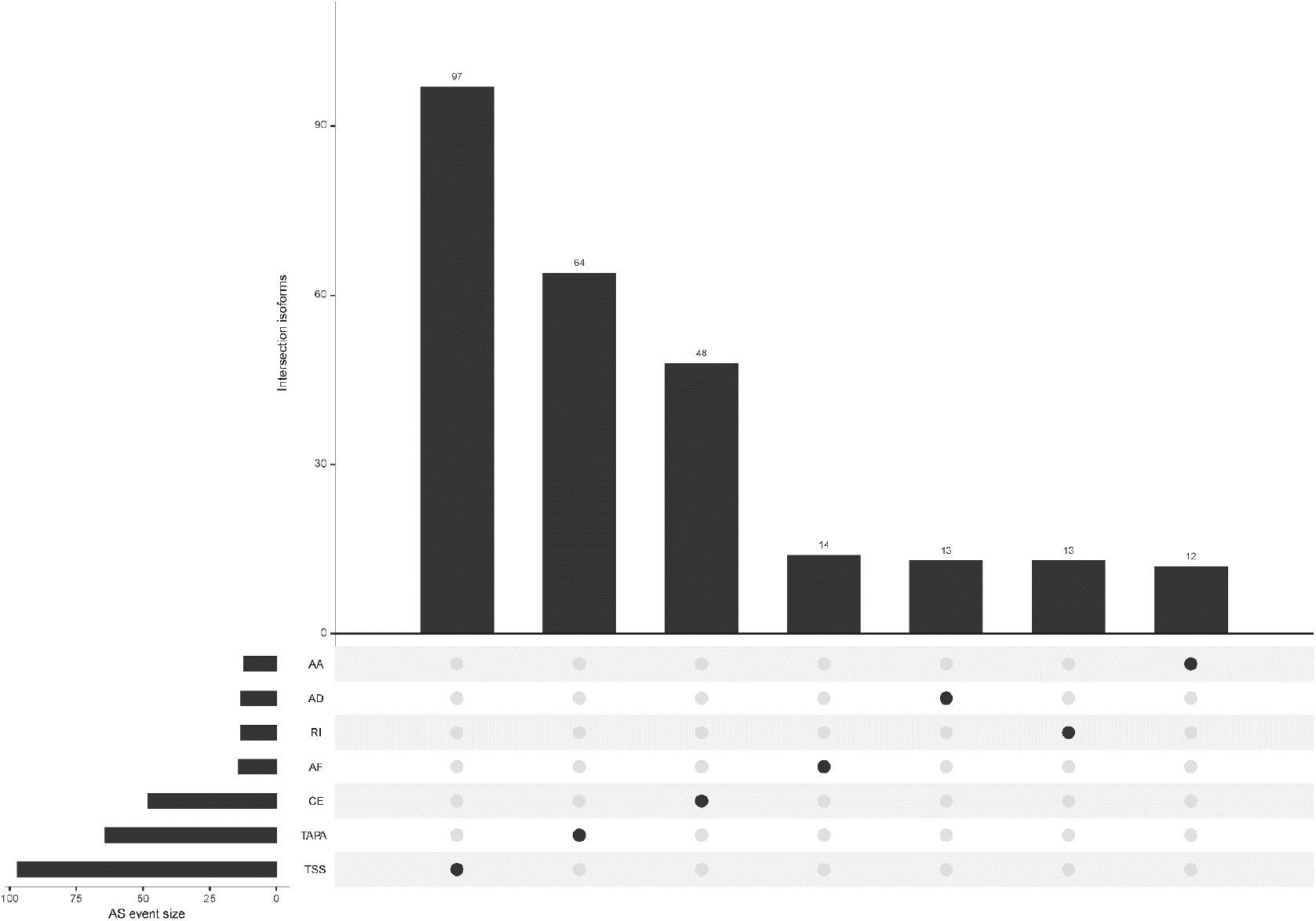
Identification of EOGC-specific AS events. (a) Heatmap of the ESAS between 80 paired tumor tissues and 79 adjacent normal tissues (absolute fold change ⩾ 0.1, probability > 0.9); (B) The landscape of 267 ESAS shown in an UpSet plot.

For the single gene with multiple ESAS events, the regulatory directions of varied ESAS events between tumor and normal tissues can be either the same or opposite. Eight genes (COL5A1, CBWD1, SLC47A2, SSPO, AL020989.1, RPL5P1, AACSP1, and AC022905.1) exhibited the same regulatory direction of AS events; while the other 22 genes showed opposite regulatory direction (Table S2). Interestingly, the proportions of certain AS patterns between the ESAS and entire AS events were inconsistent. TAPA, a top pattern among all AS events (54.0%), only contributed to 24.0% of the ESAS events.

### Profile of EOGC-specific genes and gene isoforms

Based on the above results, we further studied how the dysregulated AS events affected the expressions of genes and gene isoforms. We identified 4809 genes from a total of 16383 genes, and also identified 6152 gene isoforms from a total of 16283 gene isoforms (CPM > 0.5 and the percentage of samples with read count values > 95%).

To generate EOGC-specific genes and gene isoforms, we compared the read count value of genes or gene isoforms from the tumor and normal or adjacent non-tumor tissues by limma-voom [30]. After refining, 373 genes and 469 gene isoforms were identified as EOGC-specific genes and gene isoforms, respectively (absolute fold change ⩾ 1 and FDR < 0.05, Figure 3a & 3b, Table S3). In addition, we profiled the enriched TF binding motifs in promoters of the EOGC-specific genes and gene isoforms (Table S4). A total of four genes and three gene isoforms with ESASs were differentially expressed in the EOGC (Figure 3c).

**Figure 3.**
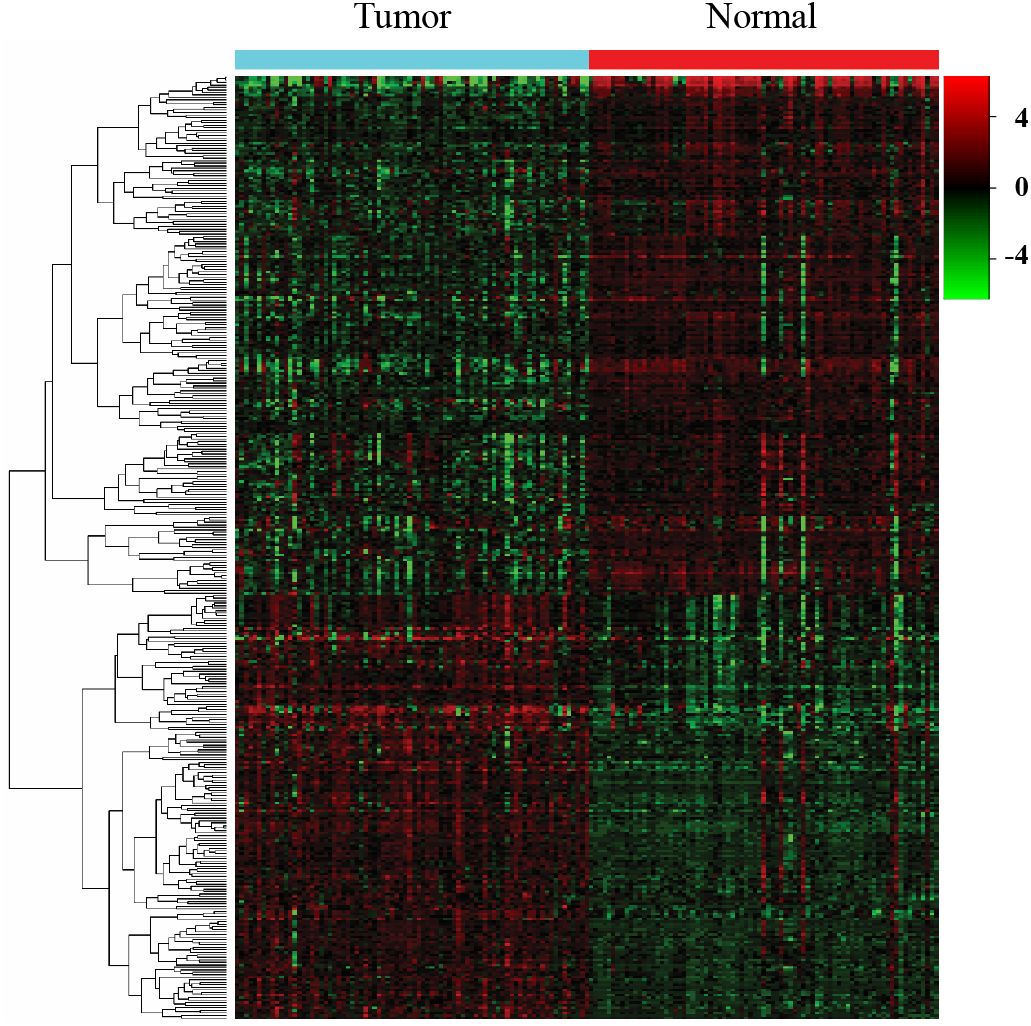
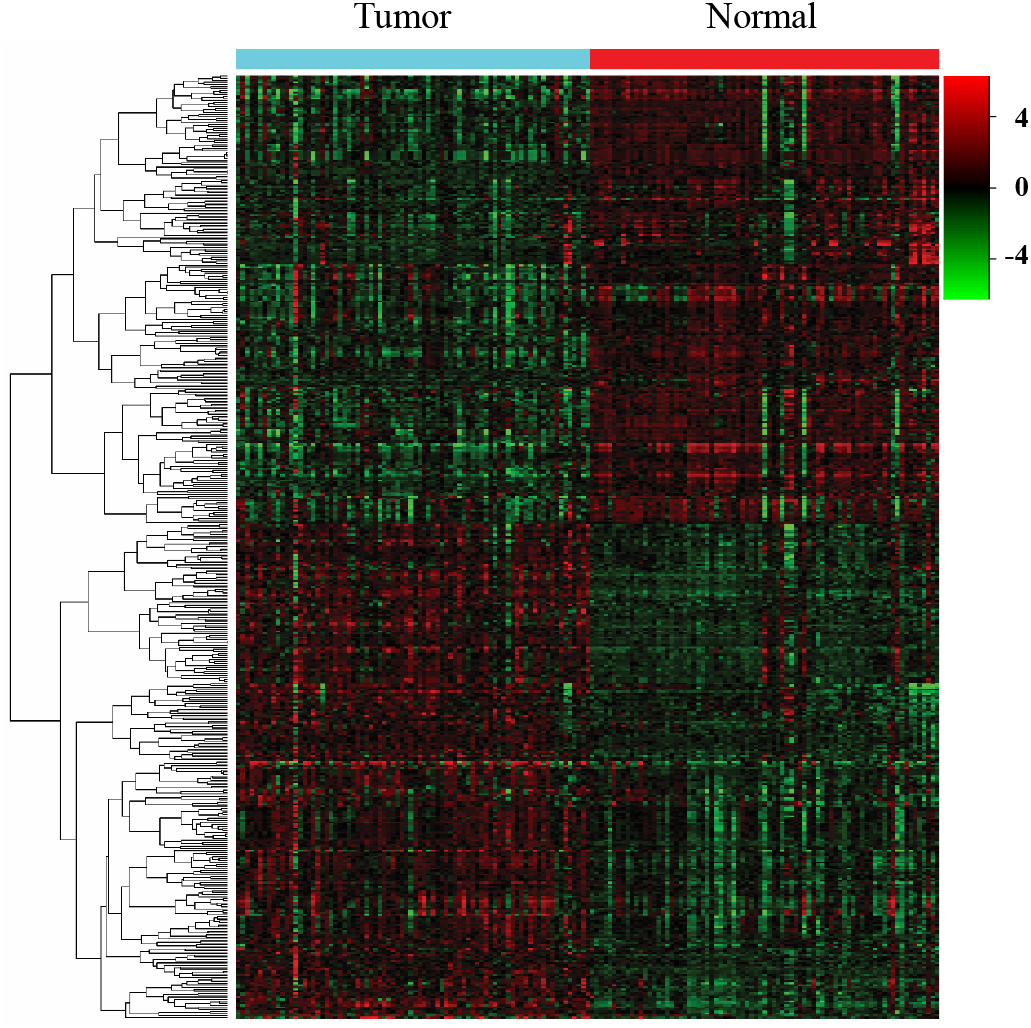
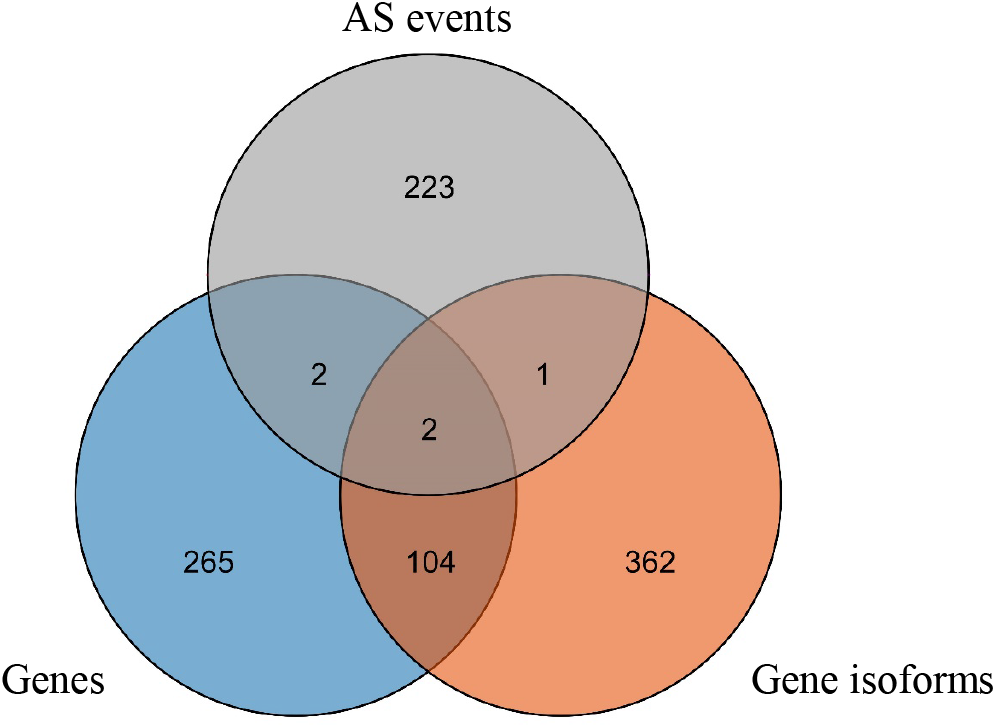
Identification of EOGC-specific genes and gene isoforms. (a) Heatmap of the EOGC-specific genes between 80 paired tumor tissues and 79 adjacent normal tissues (absolute fold change ⩾ 1, FDR < 0.05); (b) Heatmap of the EOGC-specific gene isoforms between 80 paired tumor tissues and 79 adjacent normal tissues (absolute fold change ⩾ 1, FDR < 0.05); (c) the illustration of intersection set of ESAS, EOGC-specific genes and gene isoforms by Venn diagram.

### Pathway and protein-protein interaction (PPI) network of ESAS events and EOGC-specific genes and gene isoforms

To understand the biological roles of the ESAS and EOGC-specific genes and gene isoforms, the GO and KEGG pathway analyses were performed. The results showed that the ESAS parent genes were linked to the GO terms of carboxylic acid biosynthetic process (GO:0046394), organic acid biosynthetic process (GO:0016053), purine-containing compound metabolic process (GO:0072521), collagen-containing extracellular matrix (GO:0062023), etc.. These ESAS parent genes were also found to be enriched in KEGG signaling pathways, including Nicotine addiction (hsa05033) and Arginine and proline metabolism (hsa00330) (Figure 4a and Table S5). The GO and KEGG pathway results of the EOGC-specific genes and gene isoforms were also included in the supplementary table (Table S5).

**Figure 4.**
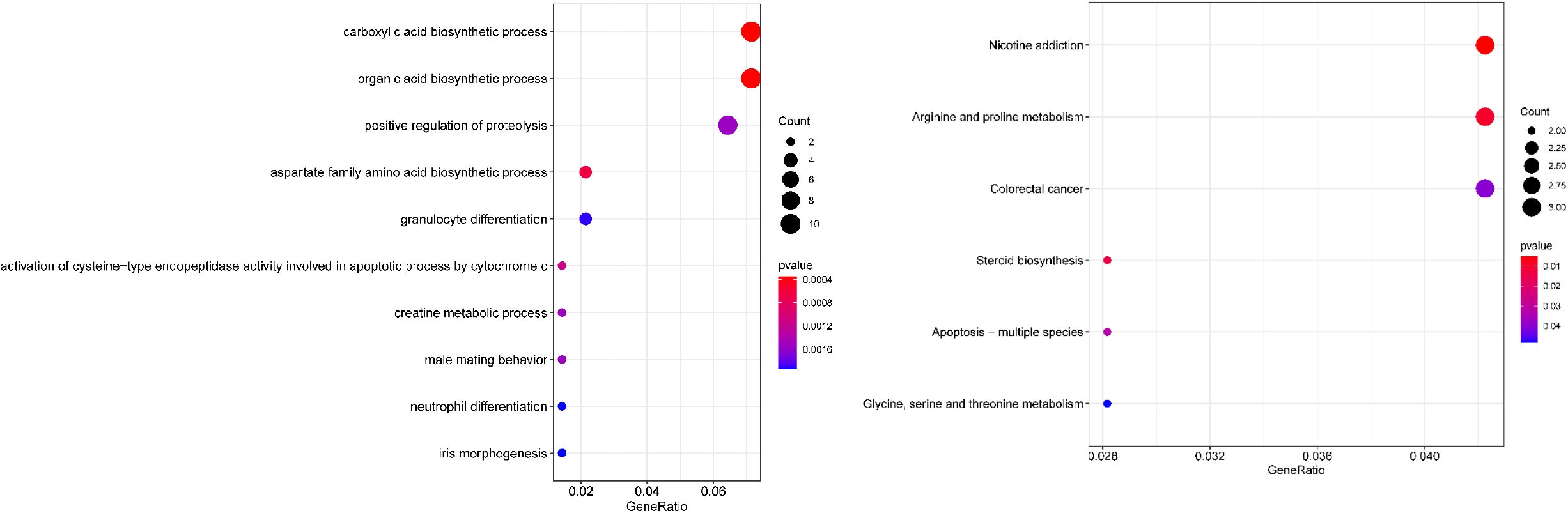
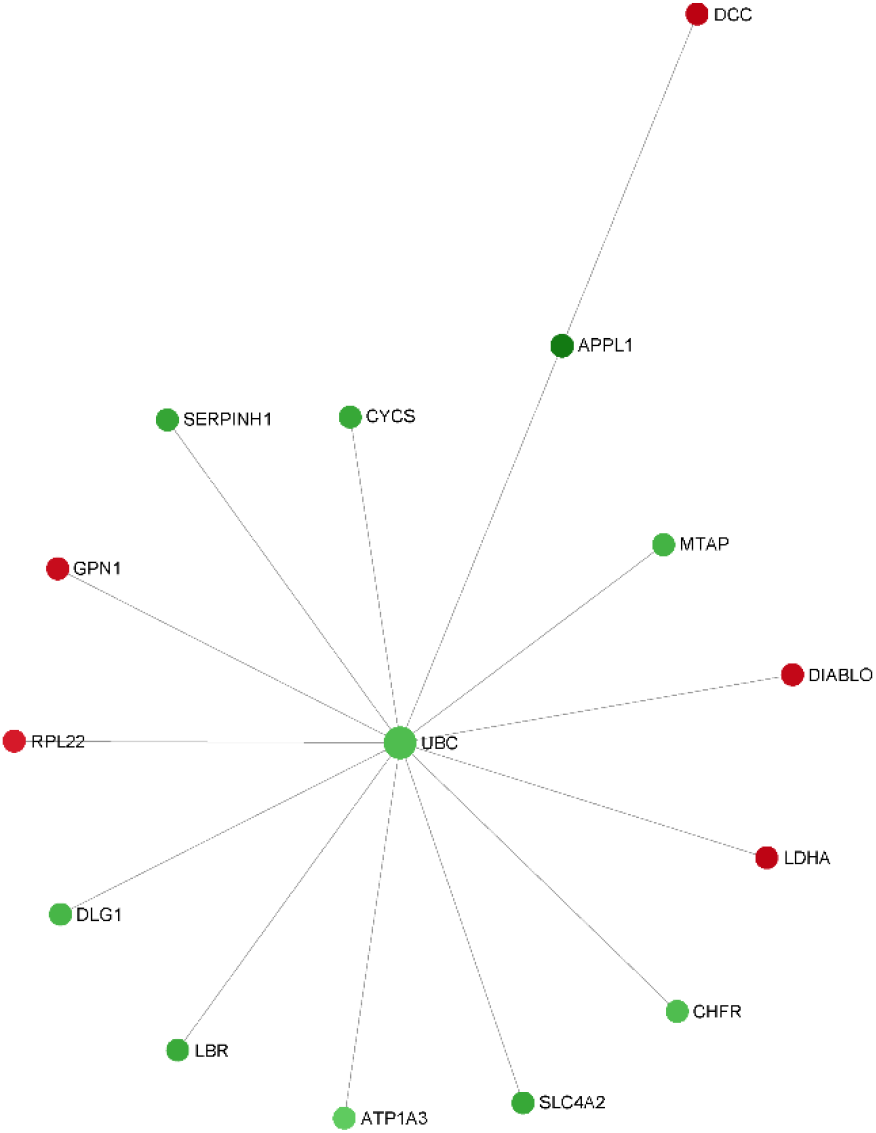
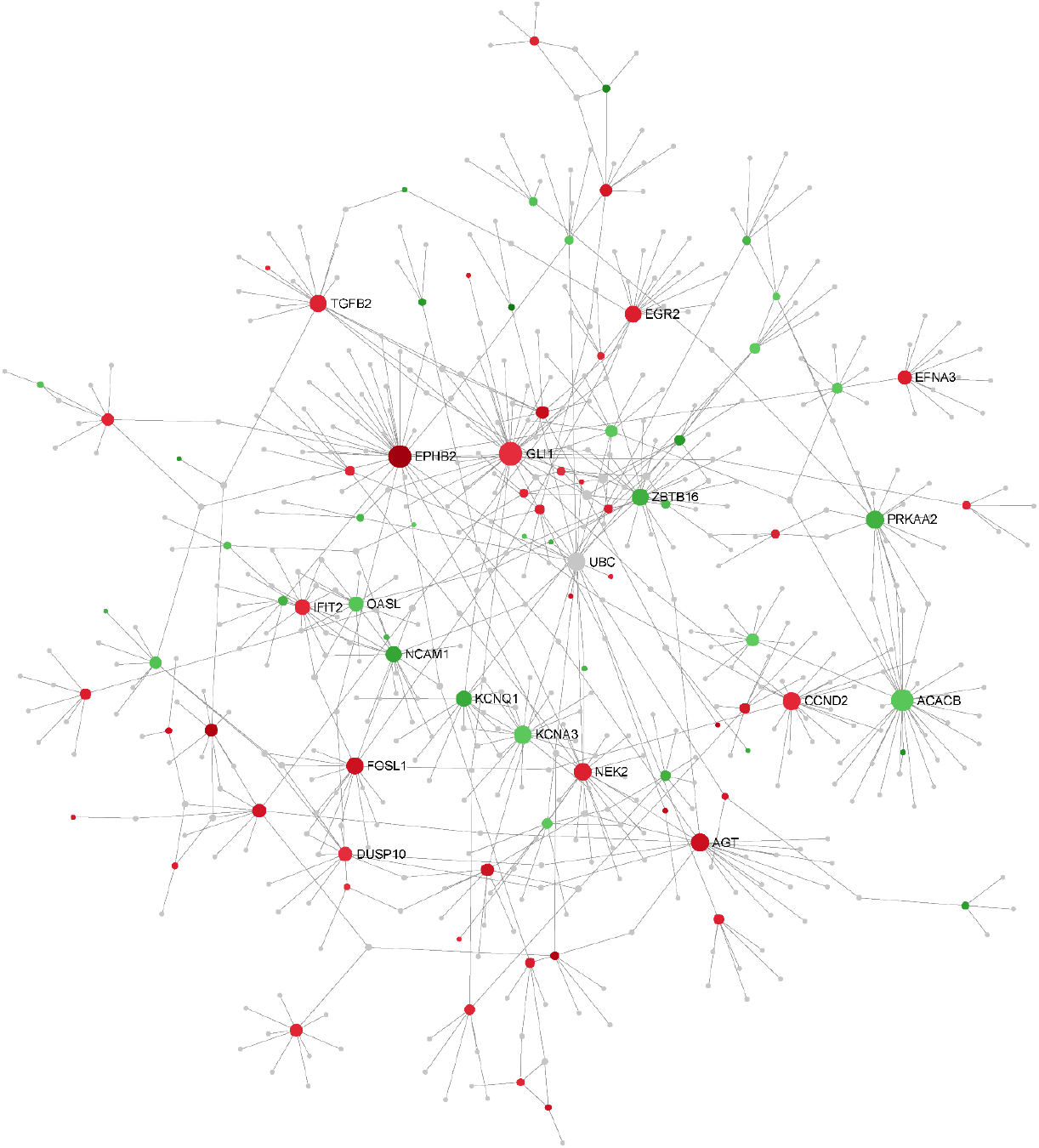
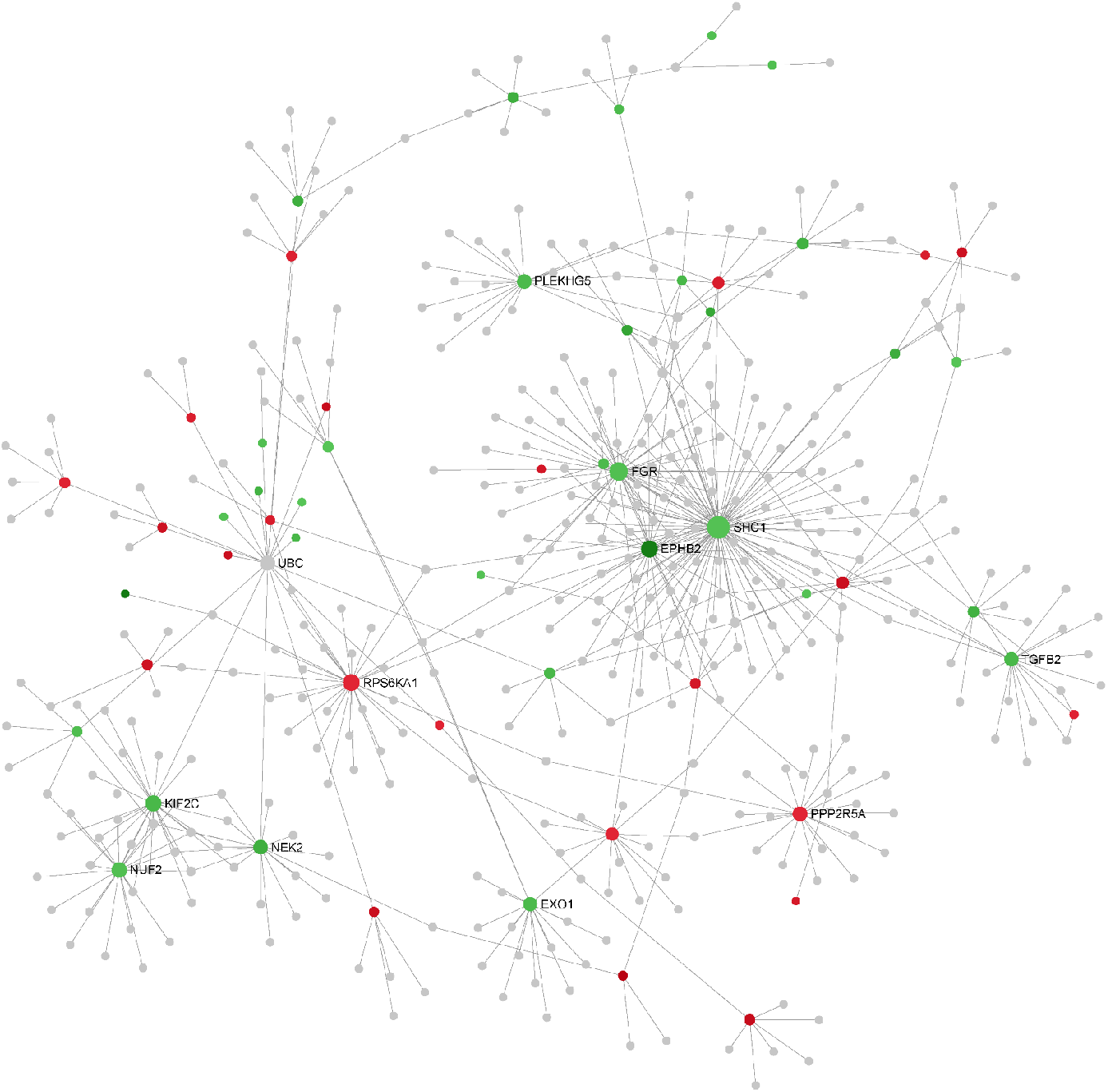
Pathway and protein-protein interaction (PPI) network of ESAS events, EOGC-specific genes, and gene isoforms. (a) Top 10 GO and KEGG pathways with P-value < 0.05; (b) The protein-protein interaction network of ESAS events (Zero-order network); (c & d) The protein-protein interaction network of EOGC-specific genes and gene isoforms, respectively (first-order network). Red color: up-regulation; Green color: down-regulation.

The pathway analysis results showed that the parent genes of ESAS might play a vital role in regulating the cancer-related biological processes. The parent genes of ESAS were further analyzed by the PPI network. Using the Zero-order network, we built the PPI network of the ESAS parent genes, which contained 15 nodes, 14 edges and 15 seeds (Figure 4b, Table S6). We found that UBC was the hub gene in the network. Meanwhile, based on the first-order network, we also built 29 and 27 PPI networks using EOGC-specific genes and gene isoforms, respectively (Figure 4c&4d, Table S6). UBC, NEK2, EPHB2, and DCTN1 genes were identified as the hub genes in two or more of these networks (DCTN1 was identified as hub gene in the section of protein modification analysis).

### Relationship between ESAS and immune cell infiltration

To untangle the relationship between the hub genes and gastric tumorigenesis, we analyzed the hub genes in TCGA STAD dataset using TIMER2 database. We found that UBC, NEK2, EPHB2, and DCTN1 genes were dysregulated in TCGA STAD dataset (p-value <0.001, Figure 5a). The correlation between the expression of the hub genes and cell abundances in STAD dataset was analyzed. We found that the expression of the UBC gene was positively correlated with the infiltration level of common lymphoid progenitor cells but was negatively correlated with the infiltration level of neutrophil cells. The expression of NEK2 gene was positively correlated with the infiltration level of myeloid-derived suppressor cells (MDSC) but was negatively correlated with the infiltration level of hematopoietic stem cells. The expression of EPHB2 gene was positively correlated with the infiltration level of NK cells but was negatively correlated with the infiltration level of activated myeloid dendritic cells. The expression of DCTN1 gene was positively correlated with the infiltration level of endothelial cells but was negatively correlated with the infiltration level of common lymphoid progenitor cells (Figure 5b, Table S7). The above findings supported our conclusion that the hub genes played vital roles in gastric tumorigenesis and indicated the potential relationships between the AS events and cell abundance.

**Figure 5.**
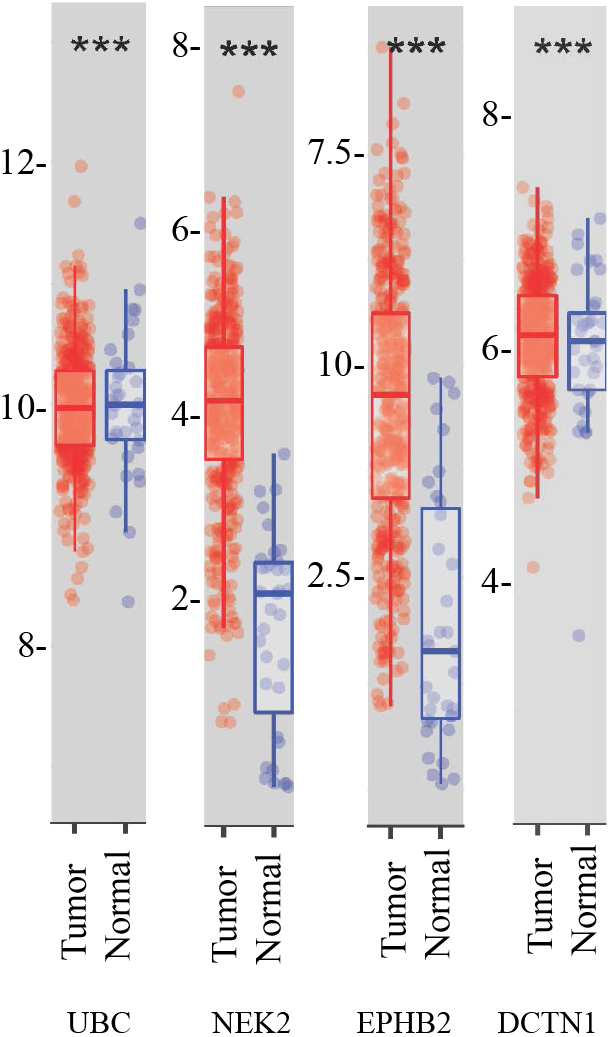
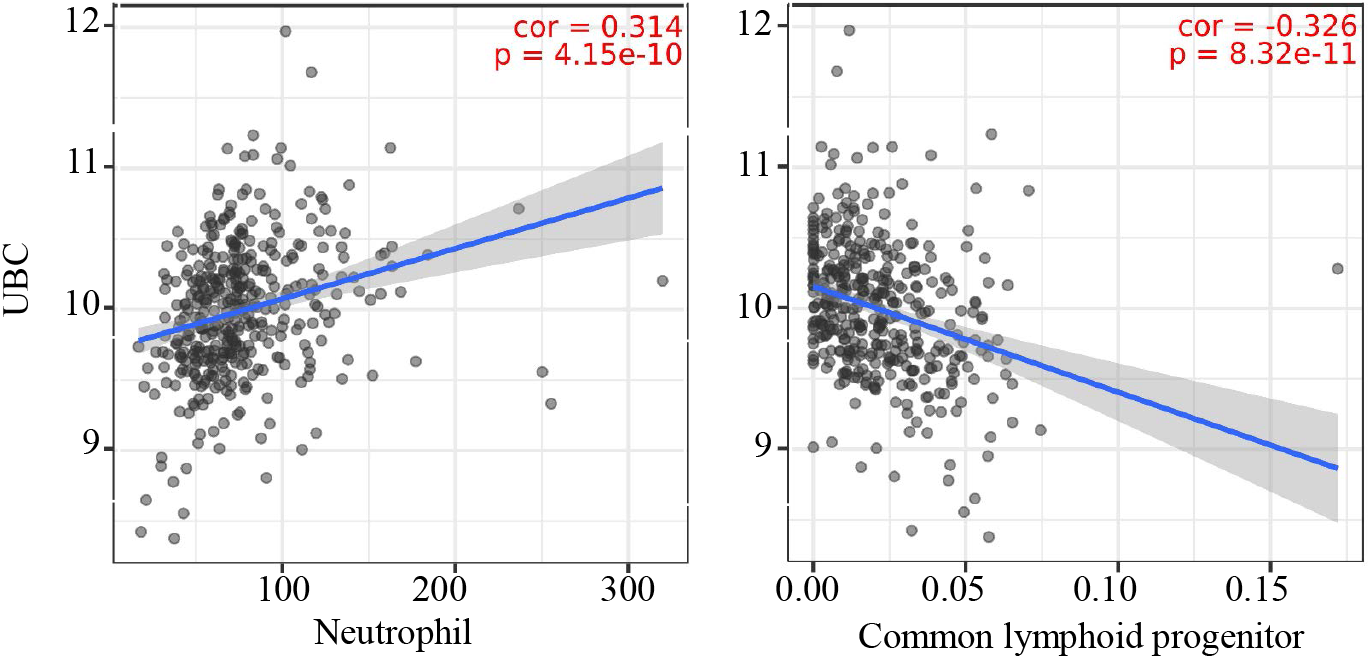
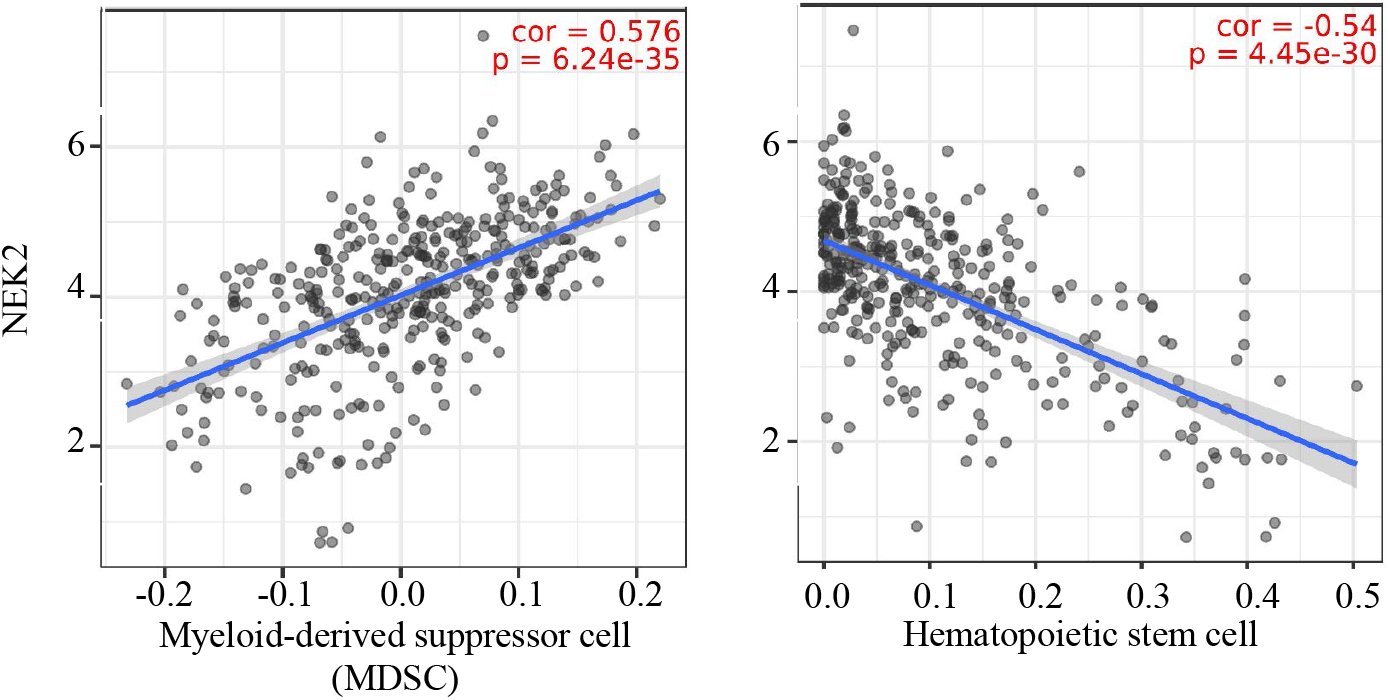
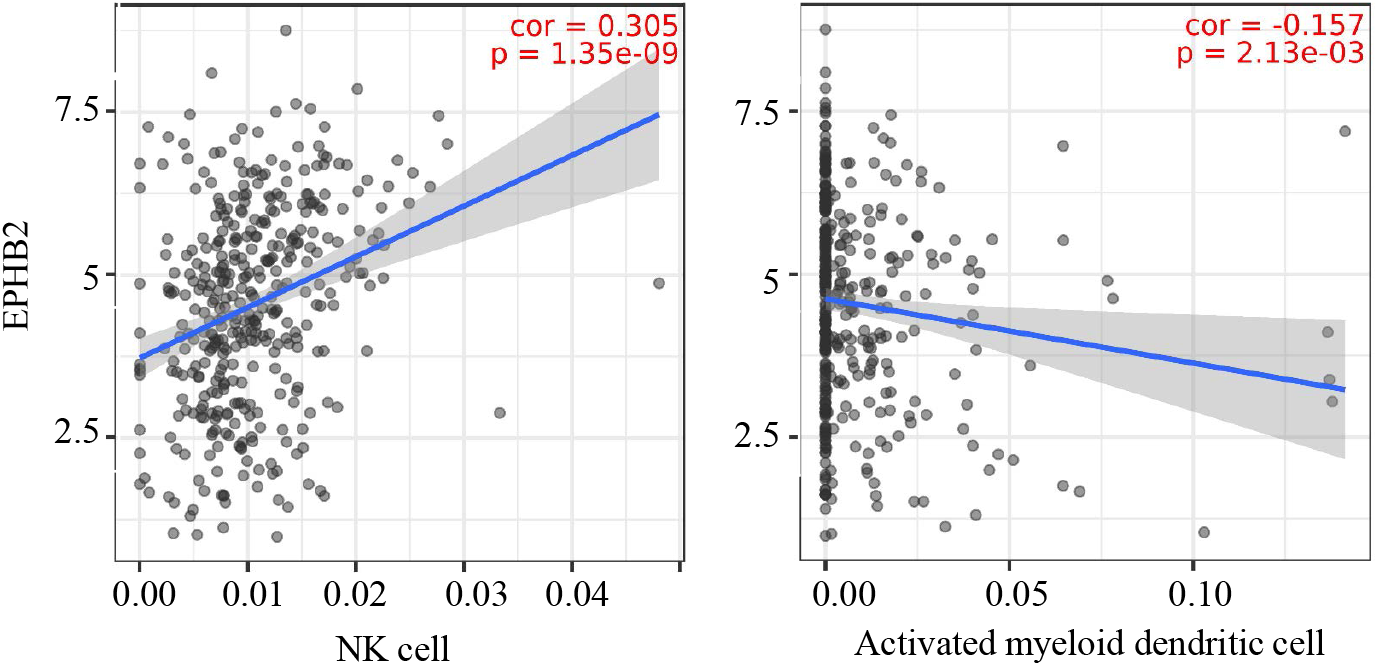
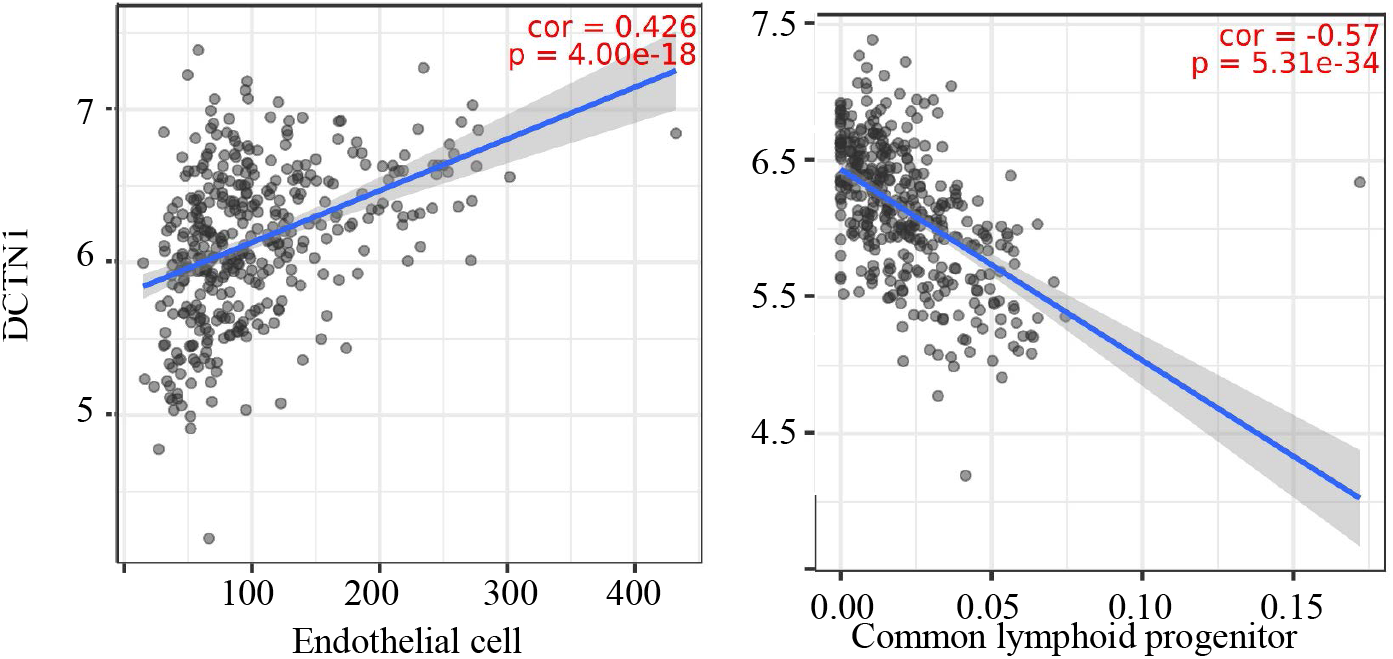
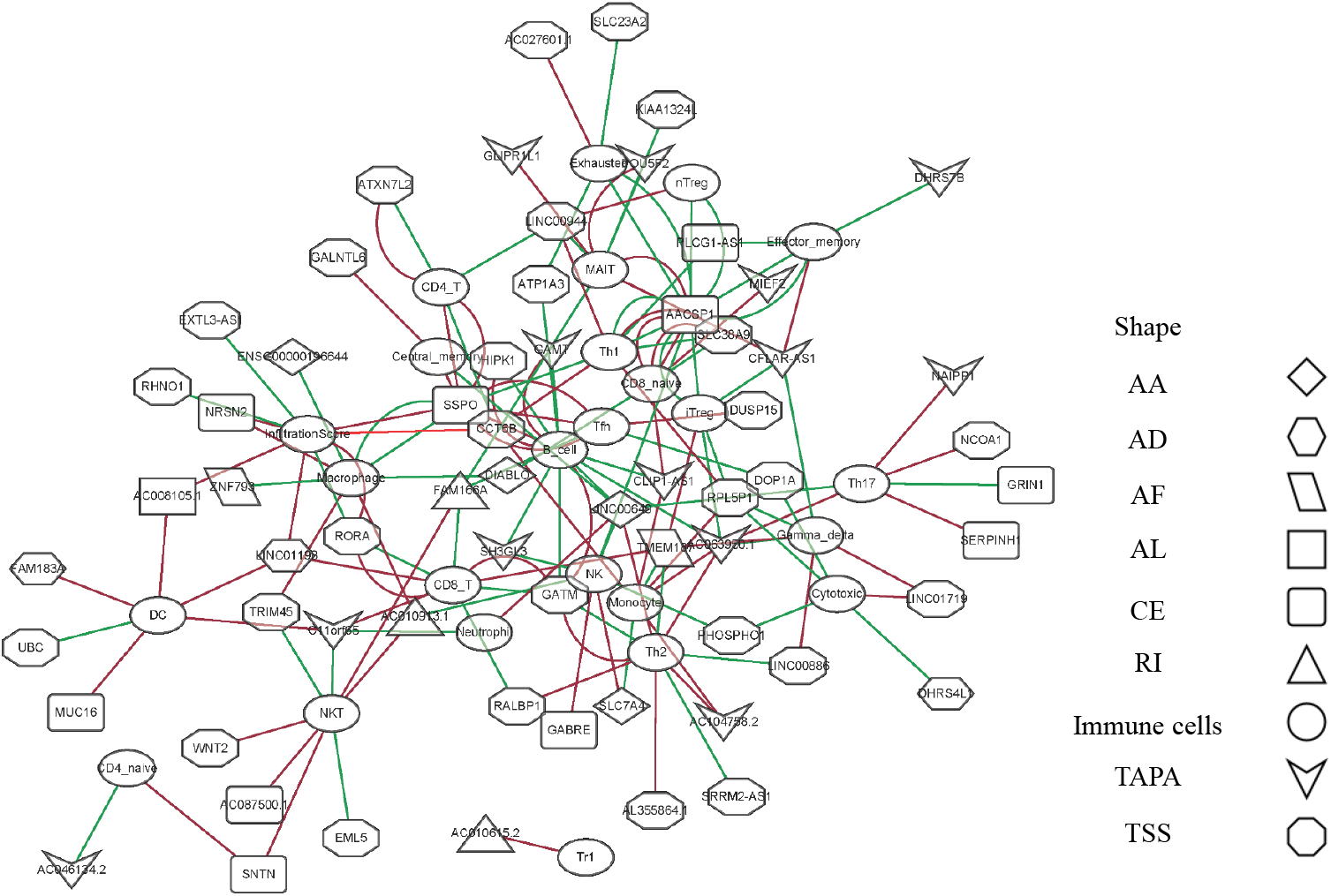
Relationship between ESAS and immune cell infiltration. (a) The expressions of UBC, NEK2, EPHB2, and DCTN1 in TCGA STAD cohort(***: p-value <0.001); (b) The representative plots of the immune infiltration and the expressions of UBC, NEK2, EPHB2, and DCTN1 in TCGA STAD cohort; (c) The Spearman’s rank correlation network of ESAS and 24 types of immune cells (including the infiltration score). Red color: positive correlation; Green color: Negative correlation. Hub gene and cell abundance in TCGA STAD cohort

Here, we postulated that the ESAS might be also involved in the immune cell infiltration. To test the hypothesis, the immune cell abundancy in EOGC was calculated ImmuCellAI database. Then, Spearman’s rank correlation analyses were performed to indicate the relationship between the ESAS and immune cell infiltration in EOGC. In total, 24 types of immune cells (including the infiltration score) were correlated with 77 ESAS events (Spearman’s rank correlation coefficients > 0.4 or < −0.4, BH adjusted p-value < 0.05, Figure 5c, Table S8).

### Network of ESAS and regulatory factors

The AS events are regulated by various factors, including splicing factors (SFs) and APA core factors. However, it is unknown how ESAS are regulated by these factors in EOGC. To gain insights into this, we built the correlation network between the expression of 71 experimentally validated SFs, 22 APA core factors, and the PSI values of ESASs [25, 26].

In the network of ESAS and SFs, 70 ESASs were associated with 25 SFs (Figure 6a, Table S9). All SFs were significantly correlated with at least five AS events. Moreover, one AS event could be regulated by up to 28 different SFs. We also constructed a network of APA core factors and TAPAs, which included seven APA core factors and ten ESAS events, respectively (Figure 6b, Table S9).

**Figure 6.**
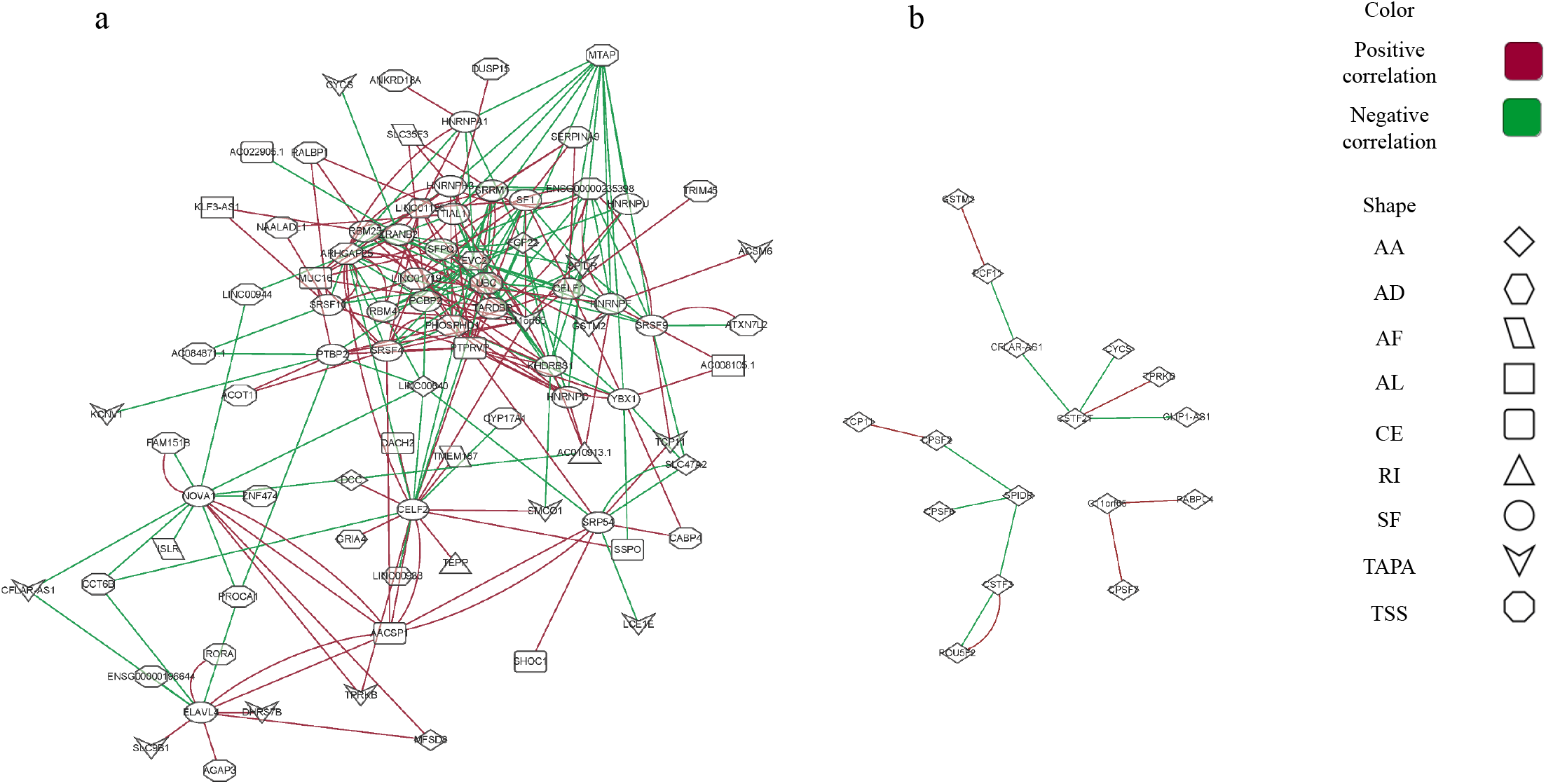
Network of ESAS and regulatory factors. (a) The Spearman’s rank correlation network of ESAS and SFs; (b) The Spearman’s rank correlation network of TAPA and APA core factors. Red color: positive correlation; Green color: Negative correlation.

### Clinically relevant and protein modification associated AS events

There are few analyses to identify clinically relevant and protein modification associated AS events in EOGC or GC. In this study, we identified 864 differentially expressed AS events from 577 genes between EBV positive and negative tumor tissues, 3335 AS events from 2056 genes associated with MSI-High (MSI-H) and MSI-Low (MSI-L) tumor tissues (absolute fold change ⩾ 1 and FDR < 0.05) (Figure 7a, Table S10).

**Figure 7.**
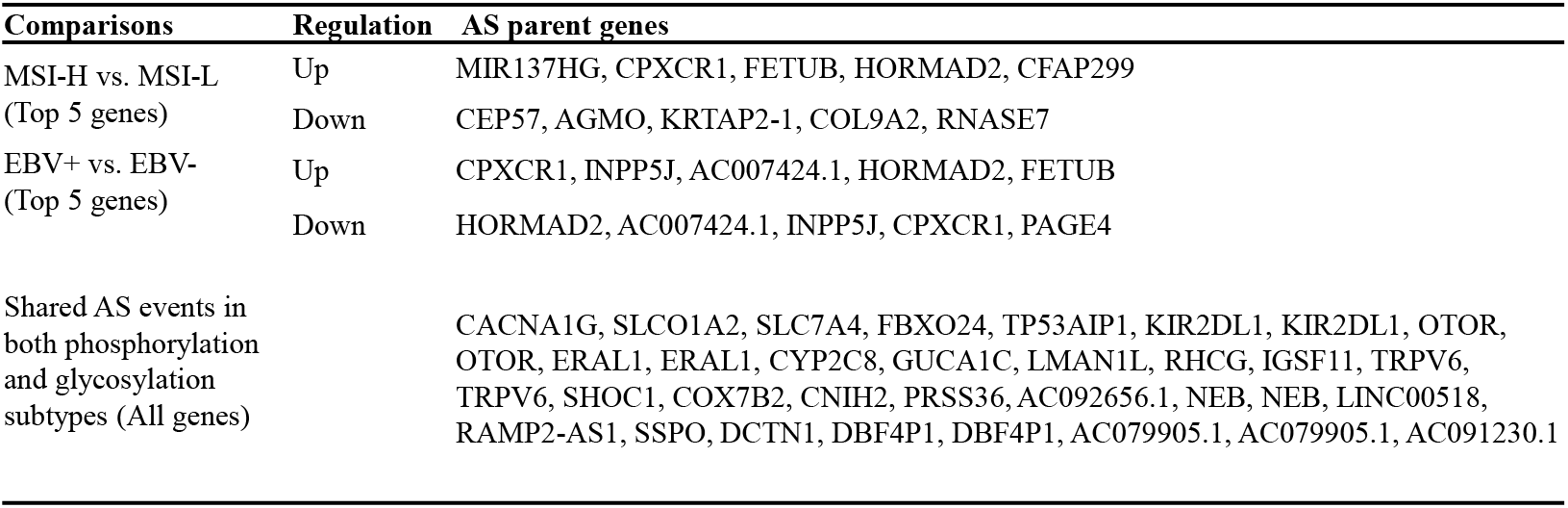
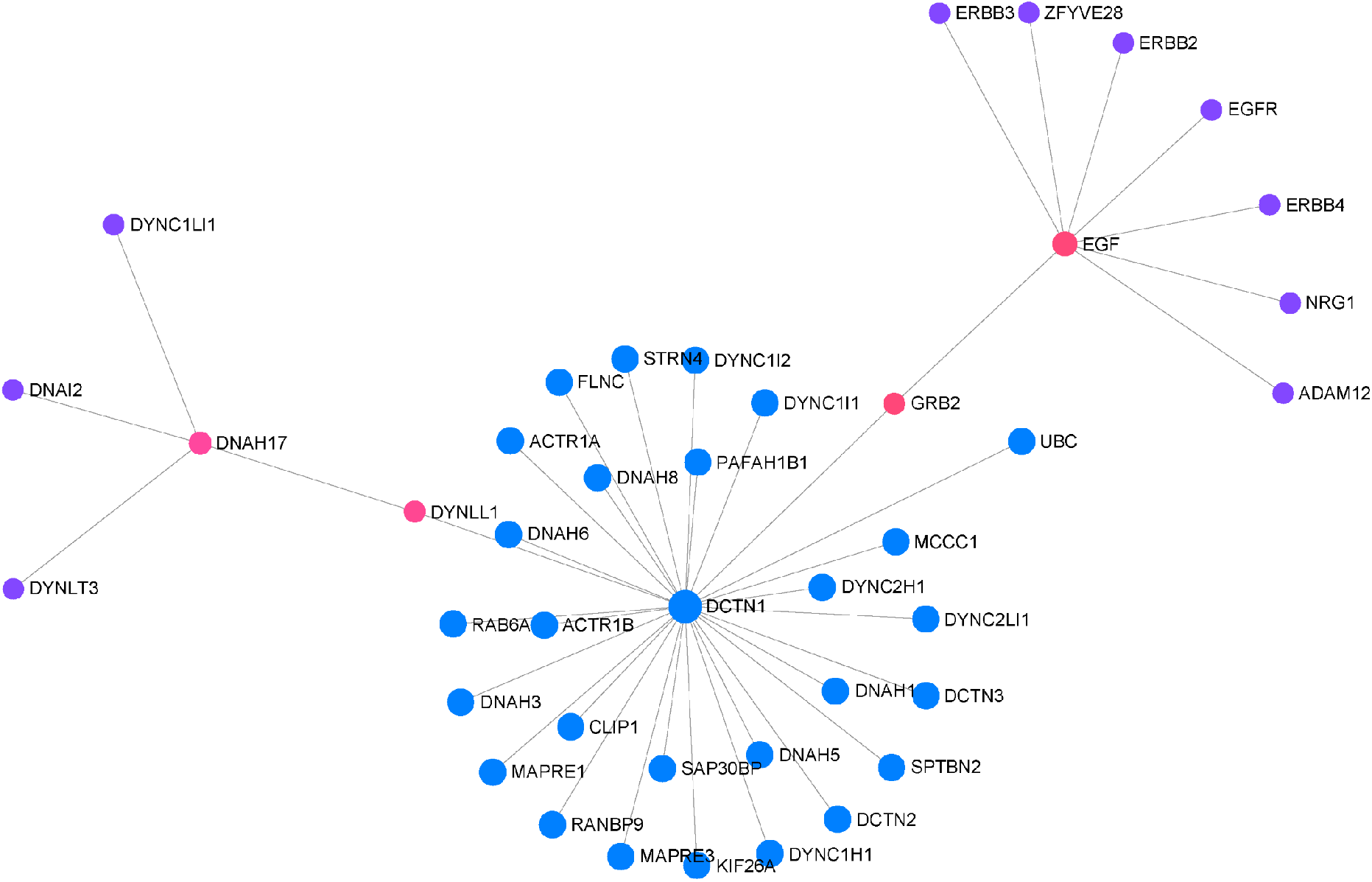
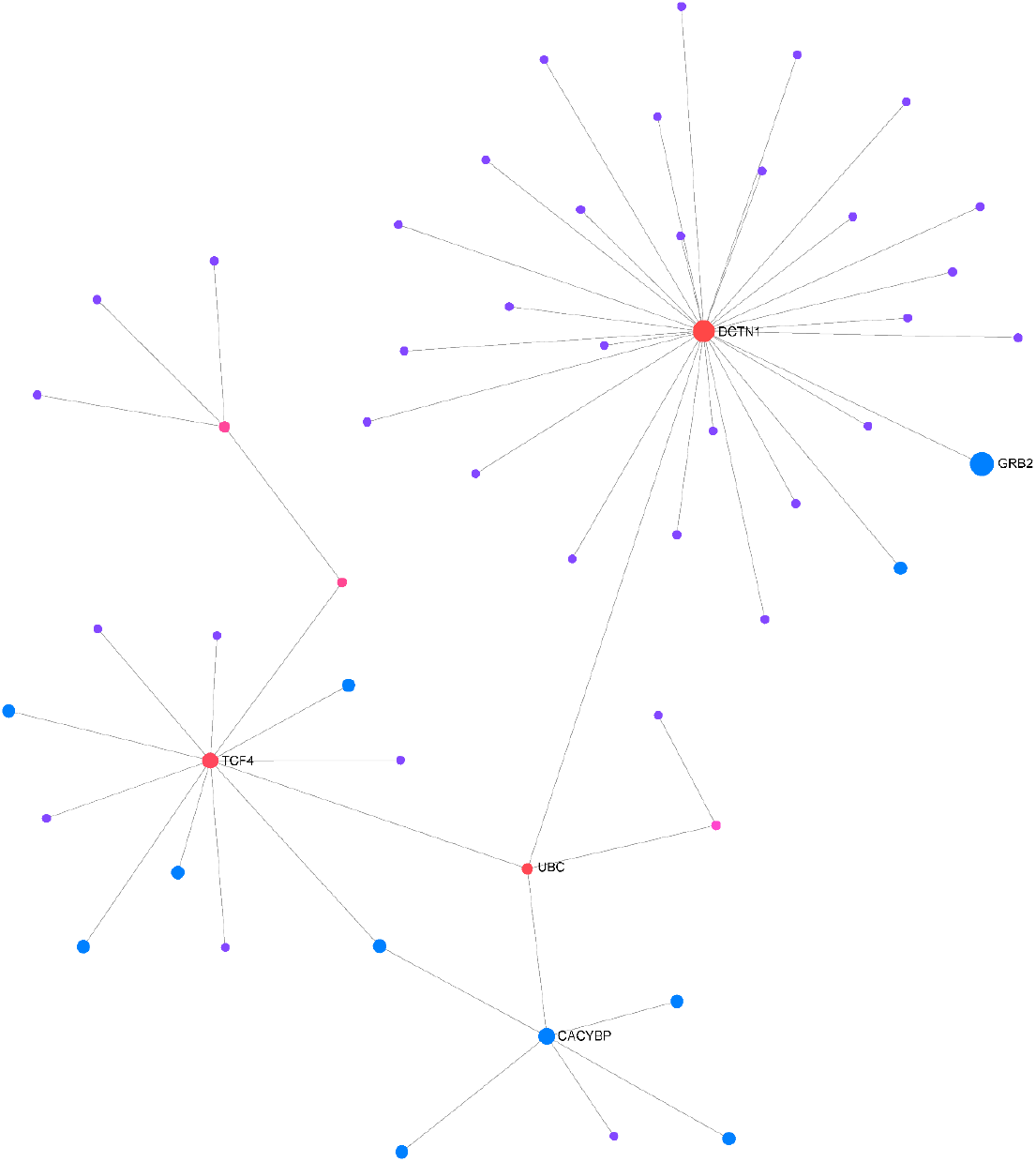
Clinically relevant and protein modification associated AS events. (a) Clinically relevant AS events; (b & c) The protein-protein interaction (first-order) networks of Glycosylation subtype and Phosphorylation subtype related AS event parent genes, respectively.

The patients were subtyped according to the status of protein modifications, including phosphorylation and glycosylation, by a previous study [5]. We found that 82 AS events from 60 genes were differentially expressed among phosphorylation subtypes one to three, while the other 85 AS events from 65 genes were differentially expressed among glycosylation subtypes one to three (absolute fold change ⩾ 1 and FDR < 0.05) (Figure 7b&7c, Table S10). In addition, a total of 27 AS events were dysregulated in both the phosphorylation and glycosylation subtypes (Figure 7a). DCTN1 gene is a key hub gene in both phosphorylation and glycosylation networks.

## Discussion

Investigation of AS events using TCGA dataset or has been a promising method and resulted in better understanding of the roles of mRNA processing in the malignancy diseases [31–33]. However, it remains challenging to reveal the roles of the AS events in certain subtypes of cancer. The unique subtype of cancer can be contributed by the AS events during the tumorigenesis. For example, the DOCK5 gene variant was identified as an oncogenic isoform in HPV-negative head and neck squamous cell carcinoma [34]. Herein, we conducted a systematic study to profile AS events in EOGC. In addition, we elucidated the roles of ESAS events with splicing factors, APA core factors, immune cell infiltration, and protein modifications.

The distribution of AS events in gastric cancer was tissue-specific and related to the prognosis of patient [35]. Furthermore, the expression patterns of AS events in gastric cancer could be altered due to Epstein-Barr virus infection [9]. In this study, the RNA-Seq data with the protein modification status for subjects obtained from a published report were analyzed [5]. In addition, we used Whippet to map the fastq format file to the reference genome, which generated two additional AS patterns (TSS and TAPA) than SpliceSeq [36] or MATS [37]. These two approaches made the results more accurate and comprehensive than studies based on TCGA SpliceSeq database [38].

According to our results, 267 ESAS, 6152 gene isoforms, and 4809 genes were aberrantly expressed in EOGC patients. The parent genes of the ESAS were significantly enriched in GO and KEGG. Further analyses indicated that the UBC, NEK2, EPHB2, and DCTN1 genes were the hub genes in PPI networks. It was reported that the UBC gene regulated the cell ubiquitin under normal and stressful conditions and the TSS events were identified within the promoter region of UBC gene [39]. The NEK2 protein was considered a splicing factor kinase through phosphorylation of the splicing factors, including the oncogenic SRSF1 protein [40]. Alternative splicing and alternative polyadenylation encoded a variant of EPHB2 gene which is a member of the EPH receptor protein-tyrosine kinase [41]. The genetic structure variability of DCTN1 gene was involved in the development of limb-girdle muscular dystrophy [42] and neuron differentiation [43]. These reports might shed light on the roles of the AS events and hub genes in gastric cancer pathogenesis.

The distribution of tumor-infiltrating lymphocytes in gastric cancer was correlated with the tumor histological type and clinical outcome [44]. On the basis of the immune infiltration data, the immunoscore, a prognostic signature, was established for prognostic predictions of gastric cancer [45, 46]. Our results showed that the hub genes and ESAS were correlated with distinct immune cell populations in EOGC. For example, TRIM45 gene encodes a protein to regulated MAPK and NF-κB pathways which may inhibit cancer cell proliferation [47]. In our study, TSS events of TRIM 45 gene were positively correlated with macrophages but negatively correlated with NK T cells. Thus, we postulated that the alternative splicing of genes might have potential roles in regulating the abundance of certain immune cells in EOGC.

Two major types of regulatory factors, SFs and APA core factors, participated in the modulation of AS events [13, 48]. Hence, we built two networks (SFs vs. ESAS and APA core factors vs. TAPA) based on the Spearman’s rank correlation. We found that the dysregulation of SFs was highly associated with the ESAS expression and the APA core factors were linked to the TAPA events, which is consistent with the findings in previous reports [49, 50]. Our results suggests that SF and APA core factor contributes to the post-transcriptional mRNA processing of EOGC.

Alternative splicing modulates protein modification such as protein phosphorylation [51] and protein glycosylation [52]. Alternative splicing might modulate the protein modification through the isoform switch and change of binding domain for N-linked glycosylation [53]. One isoform of CD44 has been shown to activate RAS through the phosphorylation of Erk [54]. In addition, the small molecule, SM08502, reduced the phosphorylation of splicing factors and generated isoforms of Wnt pathway genes which inhibited gastrointestinal tumors [55]. The reports about alternative splicing and protein glycosylation is limited. Several studies focused on CD44 variant 9 in gastric cancer [56] and variant 6 in colon cancer [57].

To our knowledge, the current study is the first to conduct a systematic analysis of alternative splicing and protein modification based on RNA sequence and mass spectrometry data in gastric cancer. The differential expressions of AS events among the subtypes of protein modification as well as EBV status and MSI status in EOGC indicated that the AS events of EOGC are subtype-specific. However, the shared AS events among different subtypes might demonstrate close associations of the subgroups defined by protein modification or viral infection in EOGC.

In conclusion, implementation of rigorous criteria ensured the identification of specific AS events related to EOGC. In total, 267 ESAS events, 6152 gene isoforms, and 4809 genes were identified and might play vital roles in EOGC tumorigenesis. The hub genes of PPI networks and ESAS events might be valuable in deciphering the immune microenvironment in early-onset gastric carcinogenesis. In addition, SF and APA core factor correlation networks revealed the underlying pathways of the splicing modulation. Furthermore, a comprehensive analysis of subtype-specific AS events suggested certain connection between protein modification and viral infection in EOGC. The findings in this study might be valuable in clinical diagnosis and prediction of early-onset gastric cancer.

## Acknowledgment

The work was partially supported by the National Cancer Institute of the National Institutes of Health (R15CA213103) and Texas A&M University T3 grant (247099).

## Notes

### Competing Interest Statement

The authors have declared no competing interest.

## References

1. Bray F, Ferlay J, Soerjomataram I, Siegel RL, Torre LA, Jemal A. Global cancer statistics 2018: GLOBOCAN estimates of incidence and mortality worldwide for 36 cancers in 185 countries. CA Cancer J Clin. 2018; 68: 394–424.

2. Sung H, Siegel RL, Rosenberg PS, Jemal A. Emerging cancer trends among young adults in the USA: analysis of a population-based cancer registry. Lancet Public Health. 2019; 4: e137–e47.

3. Takatsu Y, Hiki N, Nunobe S, Ohashi M, Honda M, Yamaguchi T, et al. Clinicopathological features of gastric cancer in young patients. Gastric Cancer. 2016; 19: 472–8.

4. Cho SY, Park JW, Liu Y, Park YS, Kim JH, Yang H, et al. Sporadic Early-Onset Diffuse Gastric Cancers Have High Frequency of Somatic CDH1 Alterations, but Low Frequency of Somatic RHOA Mutations Compared With Late-Onset Cancers. Gastroenterology. 2017; 153: 536–49 e26.

5. Mun DG, Bhin J, Kim S, Kim H, Jung JH, Jung Y, et al. Proteogenomic Characterization of Human Early-Onset Gastric Cancer. Cancer Cell. 2019; 35: 111–24 e10.

6. Modrek B, Lee C. A genomic view of alternative splicing. Nat Genet. 2002; 30: 13–9.

7. Black DL. Protein diversity from alternative splicing: a challenge for bioinformatics and post-genome biology. Cell. 2000; 103: 367–70.

8. Shi Y, Chen Z, Gao J, Wu S, Gao H, Feng G. Transcriptome-wide analysis of alternative mRNA splicing signature in the diagnosis and prognosis of stomach adenocarcinoma. Oncology reports. 2018; 40: 2014–22.

9. Armero VES, Tremblay MP, Allaire A, Boudreault S, Martenon-Brodeur C, Duval C, et al. Transcriptome-wide analysis of alternative RNA splicing events in Epstein-Barr virus-associated gastric carcinomas. PLoS One. 2017; 12: e0176880.

10. Li X, Zhang C, Gong T, Ni X, Li J, Zhan D, et al. A time-resolved multi-omic atlas of the developing mouse stomach. Nat Commun. 2018; 9: 4910.

11. Milne AN, Carvalho R, Morsink FM, Musler AR, de Leng WW, Ristimaki A, et al. Early-onset gastric cancers have a different molecular expression profile than conventional gastric cancers. Mod Pathol. 2006; 19: 564–72.

12. Jia X, Yuan S, Wang Y, Fu Y, Ge Y, Ge Y, et al. The role of alternative polyadenylation in the antiviral innate immune response. Nat Commun. 2017; 8: 14605.

13. Xiang Y, Ye Y, Lou Y, Yang Y, Cai C, Zhang Z, et al. Comprehensive Characterization of Alternative Polyadenylation in Human Cancer. J Natl Cancer Inst. 2018; 110: 379–89.

14. Fischl H, Neve J, Wang Z, Patel R, Louey A, Tian B, et al. hnRNPC regulates cancer-specific alternative cleavage and polyadenylation profiles. Nucleic Acids Res. 2019; 47: 7580–91.

15. Lai DP, Tan S, Kang YN, Wu J, Ooi HS, Chen J, et al. Genome-wide profiling of polyadenylation sites reveals a link between selective polyadenylation and cancer metastasis. Hum Mol Genet. 2015; 24: 3410–7.

16. Elkon R, Ugalde AP, Agami R. Alternative cleavage and polyadenylation: extent, regulation and function. Nat Rev Genet. 2013; 14: 496–506.

17. Sterne-Weiler T, Weatheritt RJ, Best AJ, Ha KCH, Blencowe BJ. Efficient and Accurate Quantitative Profiling of Alternative Splicing Patterns of Any Complexity on a Laptop. Mol Cell. 2018; 72: 187–200 e6.

18. Ritchie ME, Phipson B, Wu D, Hu Y, Law CW, Shi W, et al. limma powers differential expression analyses for RNA-sequencing and microarray studies. Nucleic Acids Res. 2015; 43: e47.

19. McCarthy DJ, Chen Y, Smyth GK. Differential expression analysis of multifactor RNA-Seq experiments with respect to biological variation. Nucleic Acids Res. 2012; 40: 4288–97.

20. Conway JR, Lex A, Gehlenborg N. UpSetR: an R package for the visualization of intersecting sets and their properties. Bioinformatics. 2017; 33: 2938–40.

21. Yu G, Wang LG, Han Y, He QY. clusterProfiler: an R package for comparing biological themes among gene clusters. OMICS. 2012; 16: 284–7.

22. Xia J, Gill EE, Hancock RE. NetworkAnalyst for statistical, visual and network-based meta-analysis of gene expression data. Nat Protoc. 2015; 10: 823–44.

23. Li T, Fan J, Wang B, Traugh N, Chen Q, Liu JS, et al. TIMER: A Web Server for Comprehensive Analysis of Tumor-Infiltrating Immune Cells. Cancer Res. 2017; 77: e108–e10.

24. Miao YR, Zhang Q, Lei Q, Luo M, Xie GY, Wang H, et al. ImmuCellAI: A Unique Method for Comprehensive T-Cell Subsets Abundance Prediction and its Application in Cancer Immunotherapy. Adv Sci (Weinh). 2020; 7: 1902880.

25. Giulietti M, Piva F, D’Antonio M, D’Onorio De Meo P, Paoletti D, Castrignano T, et al. SpliceAid-F: a database of human splicing factors and their RNA-binding sites. Nucleic Acids Res. 2013; 41: D125–31.

26. Shi Y, Di Giammartino DC, Taylor D, Sarkeshik A, Rice WJ, Yates JR, 3rd, et al. Molecular architecture of the human pre-mRNA 3’ processing complex. Mol Cell. 2009; 33: 365–76.

27. Shannon P, Markiel A, Ozier O, Baliga NS, Wang JT, Ramage D, et al. Cytoscape: a software environment for integrated models of biomolecular interaction networks. Genome Res. 2003; 13: 2498–504.

28. Team RC. R: A Language and Environment for Statistical Computing. R Foundation for Statistical Computing; 2020.

29. Pertea M, Shumate A, Pertea G, Varabyou A, Breitwieser FP, Chang YC, et al. CHESS: a new human gene catalog curated from thousands of large-scale RNA sequencing experiments reveals extensive transcriptional noise. Genome Biol. 2018; 19: 208.

30. Law CW, Chen Y, Shi W, Smyth GK. voom: Precision weights unlock linear model analysis tools for RNA-seq read counts. Genome Biol. 2014; 15: R29.

31. Zhang Y, Yan L, Zeng J, Zhou H, Liu H, Yu G, et al. Pan-cancer analysis of clinical relevance of alternative splicing events in 31 human cancers. Oncogene. 2019; 38: 6678–95.

32. Li Y, Sun N, Lu Z, Sun S, Huang J, Chen Z, et al. Prognostic alternative mRNA splicing signature in non-small cell lung cancer. Cancer Letters. 2017; 393: 40–51.

33. Xia Z, Donehower LA, Cooper TA, Neilson JR, Wheeler DA, Wagner EJ, et al. Dynamic analyses of alternative polyadenylation from RNA-seq reveal a 3’-UTR landscape across seven tumour types. Nat Commun. 2014; 5: 5274.

34. Liu C, Guo T, Xu G, Sakai A, Ren S, Fukusumi T, et al. Characterization of Alternative Splicing Events in HPV-Negative Head and Neck Squamous Cell Carcinoma Identifies an Oncogenic DOCK5 Variant. Clin Cancer Res. 2018; 24: 5123–32.

35. Lin P, He RQ, Ma FC, Liang L, He Y, Yang H, et al. Systematic Analysis of Survival-Associated Alternative Splicing Signatures in Gastrointestinal Pan-Adenocarcinomas. EBioMedicine. 2018; 34: 46–60.

36. Ryan MC, Cleland J, Kim R, Wong WC, Weinstein JN. SpliceSeq: a resource for analysis and visualization of RNA-Seq data on alternative splicing and its functional impacts. Bioinformatics. 2012; 28: 2385–7.

37. Shen S, Park JW, Huang J, Dittmar KA, Lu ZX, Zhou Q, et al. MATS: a Bayesian framework for flexible detection of differential alternative splicing from RNA-Seq data. Nucleic Acids Res. 2012; 40: e61.

38. Ryan M, Wong WC, Brown R, Akbani R, Su X, Broom B, et al. TCGASpliceSeq a compendium of alternative mRNA splicing in cancer. Nucleic Acids Res. 2016; 44: D1018–22.

39. Bianchi M, Giacomini E, Crinelli R, Radici L, Carloni E, Magnani M. Dynamic transcription of ubiquitin genes under basal and stressful conditions and new insights into the multiple UBC transcript variants. Gene. 2015; 573: 100–9.

40. Naro C, Barbagallo F, Chieffi P, Bourgeois CF, Paronetto MP, Sette C. The centrosomal kinase NEK2 is a novel splicing factor kinase involved in cell survival. Nucleic Acids Res. 2014; 42: 3218–27.

41. Tang XX, Pleasure DE, Brodeur GM, Ikegaki N. A variant transcript encoding an isoform of the human protein tyrosine kinase EPHB2 is generated by alternative splicing and alternative use of polyadenylation signals. Oncogene. 1998; 17: 521–6.

42. Tokito MK, Holzbaur EL. The genomic structure of DCTN1, a candidate gene for limb-girdle muscular dystrophy (LGMD2B). Biochim Biophys Acta. 1998; 1442: 432–6.

43. Song Y, Botvinnik OB, Lovci MT, Kakaradov B, Liu P, Xu JL, et al. Single-Cell Alternative Splicing Analysis with Expedition Reveals Splicing Dynamics during Neuron Differentiation. Mol Cell. 2017; 67: 148–61 e5.

44. Pernot S, Terme M, Radosevic-Robin N, Castan F, Badoual C, Marcheteau E, et al. Infiltrating and peripheral immune cell analysis in advanced gastric cancer according to the Lauren classification and its prognostic significance. Gastric Cancer. 2020; 23: 73–81.

45. Zeng D, Zhou R, Yu Y, Luo Y, Zhang J, Sun H, et al. Gene expression profiles for a prognostic immunoscore in gastric cancer. Br J Surg. 2018; 105: 1338–48.

46. Jiang Y, Zhang Q, Hu Y, Li T, Yu J, Zhao L, et al. ImmunoScore Signature: A Prognostic and Predictive Tool in Gastric Cancer. Ann Surg. 2018; 267: 504–13.

47. Shibata M, Sato T, Nukiwa R, Ariga T, Hatakeyama S. TRIM45 negatively regulates NF-kappaB-mediated transcription and suppresses cell proliferation. Biochem Biophys Res Commun. 2012; 423: 104–9.

48. Watermann DO, Tang Y, Zur Hausen A, Jager M, Stamm S, Stickeler E. Splicing factor Tra2-beta1 is specifically induced in breast cancer and regulates alternative splicing of the CD44 gene. Cancer Res. 2006; 66: 4774–80.

49. Wang R, Zheng D, Yehia G, Tian B. A compendium of conserved cleavage and polyadenylation events in mammalian genes. Genome Res. 2018; 28: 1427–41.

50. Listerman I, Sapra AK, Neugebauer KM. Cotranscriptional coupling of splicing factor recruitment and precursor messenger RNA splicing in mammalian cells. Nat Struct Mol Biol. 2006; 13: 815–22.

51. Venables JP. Aberrant and alternative splicing in cancer. Cancer Res. 2004; 64: 7647–54.

52. Bach MA, Roberts CT, Jr., Smith EP, LeRoith D. Alternative splicing produces messenger RNAs encoding insulin-like growth factor-I prohormones that are differentially glycosylated in vitro. Mol Endocrinol. 1990; 4: 899–904.

53. Reale MA, Hu G, Zafar AI, Getzenberg RH, Levine SM, Fearon ER. Expression and Alternative Splicing of the Deleted in Colorectal Cancer (*DCC*) Gene in Normal and Malignant Tissues. Cancer Research. 1994; 54: 4493.

54. Cheng C, Yaffe MB, Sharp PA. A positive feedback loop couples Ras activation and CD44 alternative splicing. Genes Dev. 2006; 20: 1715–20.

55. Tam BY, Chiu K, Chung H, Bossard C, Nguyen JD, Creger E, et al. The CLK inhibitor SM08502 induces anti-tumor activity and reduces Wnt pathway gene expression in gastrointestinal cancer models. Cancer Lett. 2020; 473: 186–97.

56. Moreira IB, Pinto F, Gomes C, Campos D, Reis CA. Impact of Truncated O-glycans in Gastric-Cancer-Associated CD44v9 Detection. Cells. 2020; 9.

57. Indinnimeo M, Cicchini C, Giarnieri E, Stazi A, Mingazzini PL, Stipa V. Evaluation of CD44 variant 6 expression and clinicopathological factors in pulmonary metastases from colon carcinoma. Oncology reports. 2003; 10: 1875–7.

